# Dopaminergic modulation of reward discounting in healthy rats: a systematic review and meta-analysis

**DOI:** 10.1101/2020.04.03.024364

**Authors:** Jaime J. Castrellon, James Meade, Lucy Greenwald, Katlyn Hurst, Gregory R. Samanez-Larkin

**Affiliations:** Department of Psychology & Neuroscience, Duke University, Durham, NC, U.S.A; Center for Cognitive Neuroscience, Duke University, Durham, NC, U.S.A; Department of Chemistry, Duke University, Durham, NC, U.S.A; Department of Biology, Duke University, Durham, NC, U.S.A

**Keywords:** dopamine, pharmacology, discounting, delay, probability, effort, decision making, meta-analysis

## Abstract

Although numerous studies have suggested that pharmacological alteration of the dopamine (DA) system modulates reward discounting, these studies have produced inconsistent findings. Here, we conducted a systematic review and pre-registered meta-analysis to evaluate DA drug-mediated effects on reward discounting of time, probability, and effort costs in studies of healthy rats. This produced a total of 1,343 articles to screen for inclusion/exclusion. From the literature, we identified 117 effects from approximately 1,549 individual rats. Using random-effects with maximum-likelihood estimation, we meta-analyzed placebo-controlled drug effects for (1) DA D1-like receptor agonists and (2) antagonists, (3) D2-like agonists and (4) antagonists, and (5) DA transporter-modulating drugs. Meta-analytic effects showed that DAT-modulating drugs decreased reward discounting. While D1-like and D2-like antagonists both increased discounting, agonist drugs for those receptors had no significant effect on discounting behavior. A number of these effects appear contingent on study design features like cost type, rat strain, and microinfusion location. These findings suggest a nuanced relationship between DA and discounting behavior and urge caution when drawing generalizations about the effects of pharmacologically manipulating dopamine on reward-based decision making.

## Introduction

Every day, all animals make decisions that involve weighing costs and benefits. Animals regularly devalue rewards that are relatively delayed, uncertain, or require more effort than sooner, more certain, or less effortful ones. This process is known as reward discounting. For example, people often choose to eat at restaurants because it is less effortful or time consuming than cooking a meal. In this scenario, people place a greater value on food that is immediately available or easy-to-acquire.

While most individuals discount to some degree, a range of factors influence whether one discounts rewards more steeply (stronger devaluation) or not at all. In humans, for example, income, IQ, age, smoking, and BMI have all been linked to individual differences in reward discounting (de Wit et al. 2007; Reimers et al. 2009; Seaman et al. 2016). Aside from these sociodemographic and physical health factors, discounting is often disrupted in many forms of psychopathology (Lempert et al. 2019; Amlung et al. 2019). An emergent pattern suggests that disruption to circuits involved in the neurotransmission of dopamine (DA) may account for variation in discounting behavior across specific psychopathologies that are often treated with drugs that primarily act on the dopamine system (Castrellon et al. 2019; Amlung et al. 2019).

Importantly, drugs that act on the DA system have different effects depending on their targets and action. The putative DA targets for pharmacology are presynaptic synthesis, DA transporters (DAT), and agonism or antagonism of postsynaptic D1-like or D2-like receptors. Variation in results from studies testing the effect of different DA drug effects across these sites on discounting behavior suggests the need for a quantitative comparison of experiments (Cousins et al. 1994; Wade et al. 2000; Pattij and Vanderschuren 2008; Floresco et al. 2008; Bardgett et al. 2009; St. Onge et al. 2010; Koffarnus et al. 2011; Simon et al. 2011; Pardey et al. 2013; Stopper et al. 2013; Yates et al. 2014; Sommer et al. 2014; Hosking et al. 2015; Li et al. 2015; Larkin et al. 2016). Since meta-analytic methods can evaluate studies with heterogenous features such as sample characteristics and task design, outcome measures may reflect generalizable neural and cognitive functions. Here, we conducted a systematic review and meta-analysis to evaluate DA drug-mediated effects on reward discounting in studies of healthy rats (117 effects from approximately 1,549 rats). We focused on rats because of the small number of human and non-human primate studies identified (N = 4 studies) and it remains unclear whether discounting-like behaviors generalize across species (Hayden 2016; Rosati and Hare 2016; Heilbronner 2017; Susini et al. 2020).

## Methods

### Literature Search and Study Identification

A meta-analysis was conducted following the Preferred Reporting Items for Systematic Reviews and Meta-Analysis (PRISMA) guidelines (Moher 2009). From an initial in-lab library of 34 papers on pharmacological manipulation of dopamine effects on reward discounting, we developed a database of search terms to identify additional studies. We restricted the search to the PubMed database using Medical Subject Headings (MeSH) terms that are most frequently associated with papers in the library. To identify the most frequent MeSH terms, we used the MeSH on Demand tool (https://meshb.nlm.nih.gov/MeSHonDemand) to identify terms from the abstract text of each of the 34 papers. Frequently associated terms that best described the features of the studies of interest included: “animals,” “dopamine,” “reward,” “impulsive behavior,” “choice behavior,” and “delay discounting.” The terms were then combined to search for original research examining how administration of dopaminergic drugs influence reward discounting behavior using the following PubMed search string: “Dopamine” [Mesh] AND (“reward” [Mesh] OR “delay discounting” [Mesh] OR “choice behavior” [Mesh] OR “impulsive behavior” [Mesh] OR “temporal discounting” OR “probability discounting” OR “effort discounting” OR “intertemporal choice” OR “indifference point”) AND (“drug” OR “agonist” OR “antagonist”).

We restricted the meta-analysis to original studies written in English. Studies must have included a healthy animal group (including humans, non-human primates, and rats) exposed to placebo and/or drug manipulation. Healthy animals that received lesions or other surgical manipulation prior to drug delivery were excluded unless discounting behavior was unaffected by the lesion or surgery. We further limited the analysis to studies using choice tasks or questionnaires with varying levels of temporal delays, probability, or effort expenditure. To reduce the complexity of the impact of various drugs, we limited confirmatory analyses to drugs that exhibit direct primary action on either: D1-like receptors, D2-like receptors, and DAT. D1-like and D2-like receptors describe general families of receptors that include D1Rs and D5Rs (D1-like) and D2Rs, D3Rs, and D4Rs (D2-like) (Beaulieu and Gainetdinov 2011). A number of drugs that modulate a specific receptor exhibit non-selective binding to other intra-family receptors (e.g. binding to D2Rs, D3Rs, and D4Rs) (Nichols 2010; Cho et al. 2010; Löber et al. 2011) with some exceptions including the D1/D2 antagonist flupenthixol (Wetzel et al. 1998). As a result of these similarities and to ensure that our meta-analyses are sufficiently powered, we aggregate receptor subtypes using their canonical D1-like and D2-like family grouping. Since a number of DAT-modulating drugs have similar (and in some cases preferential) affinity for the norepinephrine transporter (NET), we also report the DAT meta-analytic effect excluding such NET-preferring drugs. Drugs manipulating levels of the dopamine precursor, L-DOPA were also included with the acknowledgement that too few studies may exist to meta-analyze. We excluded studies using healthy controls identified as nicotine users or relatives of patients with Parkinson’s Disease. We also excluded studies that only tested drug effects in humans over 30 years old to reduce the influence of strong age-related declines in DA receptors. Additional studies were later excluded from analysis for reasons that would prevent reliable effect size estimation (e.g. unclear or unreported sample sizes, blurry graphs, or unreported measures of variance). All of these search methods were pre-registered on the Open Science Framework prior to the start of any research activity and all literature search materials may be viewed/downloaded at: https://osf.io/27cqw/. These steps taken to exclude studies are presented in the PRISMA flowchart in **Supplementary Figure S1.**

### Data Extraction

Effect size measures were determined using means, standard deviations, standard errors, confidence intervals, and t-statistics whenever available. Effect sizes were calculated using the ‘escalc’ function provided with the ‘metafor’ R Statistics package (Viechtbauer 2010). For studies that employed a between-subjects design, we calculated the standardized mean difference (SMD) in discounting between drug and placebo groups. Since many rat studies have small sample sizes, we used the unbiased estimator of the sampling variance for between-subject effects to account for possible non-normal distributions (Hedges 1989). For studies that employed a within-subjects (repeated-measures) design, we calculated the standardized mean change score using raw score standardization (SMCR) as this provides a less biased effect size since repeated measures may be correlated (Becker 1988). Since these correlations between drug and placebo conditions are rarely reported, they were set to r = .50 to provide a conservative calculation of the variance (Morris and DeShon 2002). To evaluate the robustness of the meta-analytic effects and validate assumptions, we compared the SMCRs assuming r = .60 and compared with effect sizes calculated using the SMD measure for all effects.

For studies that did not explicitly report these values or that used sophisticated study designs, we used a plot digitizer to determine means and standard deviations (Rohatgi, Ankit 2019).We did not include studies that provided a measure of central tendency but not a measure of variance since both measures are necessary to estimate an accurate statistic. For studies that reported effects for multiple doses of the same drug, we only extracted discounting effects from the highest dose. Studies reported different metrics of discounting including: hyperbolic discounting slope “k” parameter, impulsive choice ratio (ICR), proportion of delayed/uncertain/effortful choices, area under the curve (AUC), and indifference point (also referred to as mean adjusted delay (MAD)). Many studies did not report a single discounting parameter, but instead report the proportion of smaller or larger options at varying levels of time, probability, and effort. In these cases, to simplify comparisons across studies, we averaged the reported choice proportions across cost levels. Our primary meta-analyses pool across different cost types, however we also ran exploratory meta-regressions to compare time, probability, and effort effects wherever possible. To standardize the directionality of discounting measures, effect sizes were multiplied by either 1 or −1 to ensure that positive values reflect higher discounting (e.g. higher “k”, lower AUC).

### Random-effects Meta-analysis

Meta-analytic effects were derived using the metafor R package (Viechtbauer 2010) using random effects with restricted maximum likelihood to help account for between-study variance. Specifically, we ran five confirmatory models testing the effects of: 1.) D1-like agonists, 2.) D1-like antagonists, 3.) D2-like agonists, 4.) D2-like antagonists, and 5.) DAT-modulating drugs. We used the Q-statistic to test the null hypothesis that the common effect size is zero and *I*^2^ values to assess significance due to variance explained by heterogeneity of the effects (Borenstein et al. 2011). We evaluated publication bias and study precision asymmetry with visual inspection of a funnel plot and Egger’s test (*p* < 0.05). Finally, to evaluate sensitivity of our effects to outliers, we used the permutation tool ‘leave1out’ to repeatedly sample our mixed-effects model with one effect removed.

### Exploratory Meta-analyses

Exploratory meta-regressions examined potential effects of rat strain, discounting cost types, drug delivery location, drug dose, and D2:D3 receptor affinity. For details, see Supplementary Information.

## Results

### Studies Identified and Data Extraction

The literature search, which was run on January 8, 2018, revealed 1,343 articles. After evaluation of exclusion criteria, 42 unique articles with 121 effects published between 1994 and 2017 remained for quantitative analyses (**see Supplementary Figure S1** for a flow chart). Data were extracted using a plot digitizer for nearly all studies as insufficient statistical reporting prevented reliable estimation of effect sizes and variances. The number of effects for each drug type were: D1-like agonist (k = 7), D1-like antagonist (k = 17), D2-like agonist (k = 18), D2-like antagonist (k = 45), and DAT-modulating (k = 33). As expected, only one effect size could be extracted for drugs acting on presynaptic DA—too few to analyze. Nearly all effect sizes came from within-subjects designs (k = 115) with only 6 using a between-subjects design. It should be noted that several studies reported more than one effect as a result of repeated exposure to multiple drugs or multiple samples separately exposed to different drugs. From the included studies, the majority of effect sizes were from rats (k = 117), with only a few effects in humans (k = 3) and a single effect in non-human primates (rhesus macaques). The number of effects representing discounting of varying costs were somewhat equally distributed between time (k = 46), probability (k = 43), and effort (k =32) discounting. The most common measure of discounting was the proportion of larger options chosen (k = 91), followed by the indifference point (k = 25), hyperbolic ‘k’ value (k = 3), and proportion of smaller options chosen (k = 2). Since there were too few effects in primates (k = 4), these effects were not included with rat effects in analyses but their data are provided on OSF (https://osf.io/27cqw/). All reported effects reflect acute exposure to drug. **See Tables S1-5 for details about each effect size included in quantitative meta-analyses.**

### Random-effects meta-analysis – confirmatory analyses

#### D1-like agonists

A meta-analysis across D1-like agonists did not identify a significant common effect of drug over placebo on discounting (Q_Heterogeneity_ = 5.26, p = .511, *I*^2^ < 0.00%; Cohen’s *d* = .136, SE = .118, 95% CI [−.095, .368], p = .249. Egger’s test for plot asymmetry did not suggest the presence of publication bias (z = .169, p = .865). Leave-one-out analysis suggested that removal of any one study had no impact on the overall meta-analytic effect. **See forest plot on Figure 1 and funnel plot on Supplementary Figure S2.**

**Figure 1.**
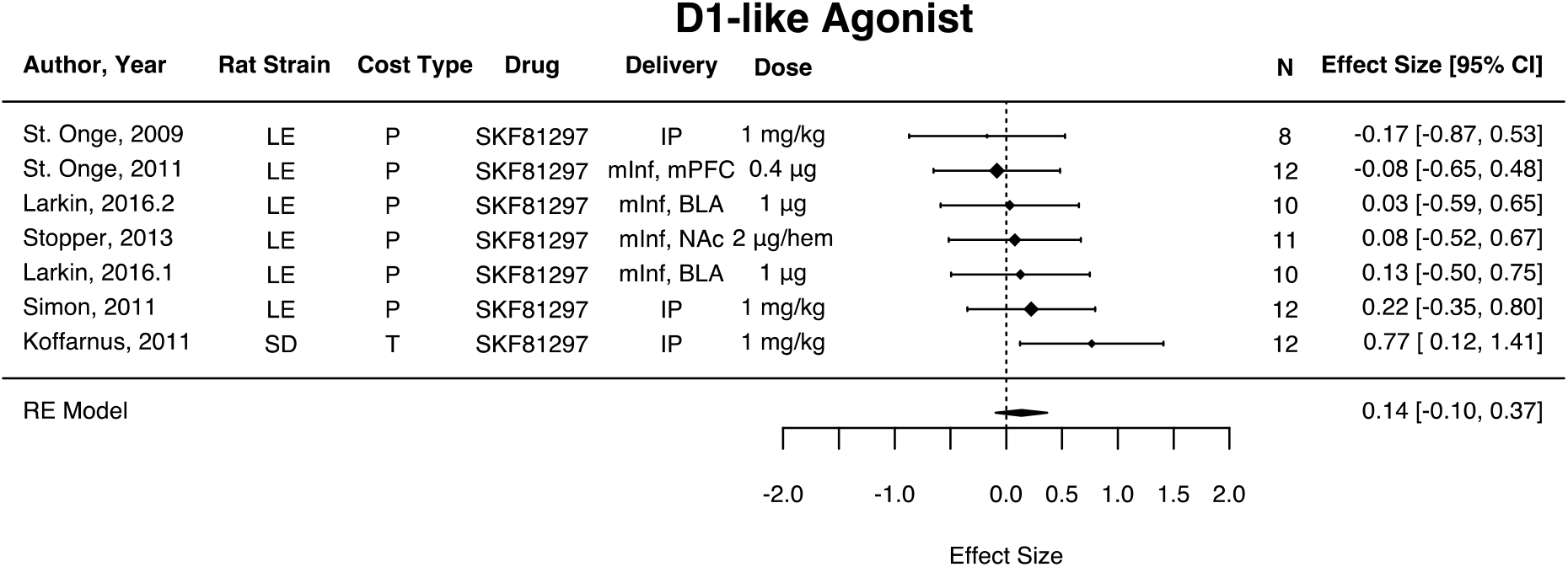
Forest plot of placebo-controlled effect of D1-like agonism on reward discounting. Positive values indicate increased discounting on drug. LE = Long-Evans, SD = Sprague-Dawley, P = Probability, T = Time, IP = intraperitoneal, mInf = microinfusion, hem = hemisphere, mPFC = medial prefrontal cortex, NAc = nucleus accumbens, BLA = basolateral amygdala.

#### D1-like antagonists

A meta-analysis across D1-like antagonists identified a significant common effect of drug over placebo on discounting (Q_Heterogeneity_ = 47.9, p < .001, *I*^2^ = 55.6%; Cohen’s *d* = .532, SE = .120, 95% CI [.296, .767], p < .001). Across effects, D1-like antagonists increased discounting over placebo. Egger’s test for plot asymmetry suggested the presence of publication bias (z = 6.11, p < .001). Leave-one-out analysis suggested that removal of any one study had no impact on the overall meta-analytic effect. **See forest plot on Figure 2 and funnel plot on Supplementary Figure S3.** Although we have chosen to group effect sizes associated with flupenthixol (a drug that antagonizes both D1Rs and D2Rs) with D2-like antagonists as reported in the next section, grouping it instead with D1-like antagonists slightly decreases but does not meaningfully alter the effects of D1-like antagonists on discounting (Q_Heterogeneity_ = 85.2, p < .001, *I*^2^ = 50.8%; Cohen’s *d* = .436, SE = .078, 95% CI [.283, .589], p < .001).

**Figure 2.**
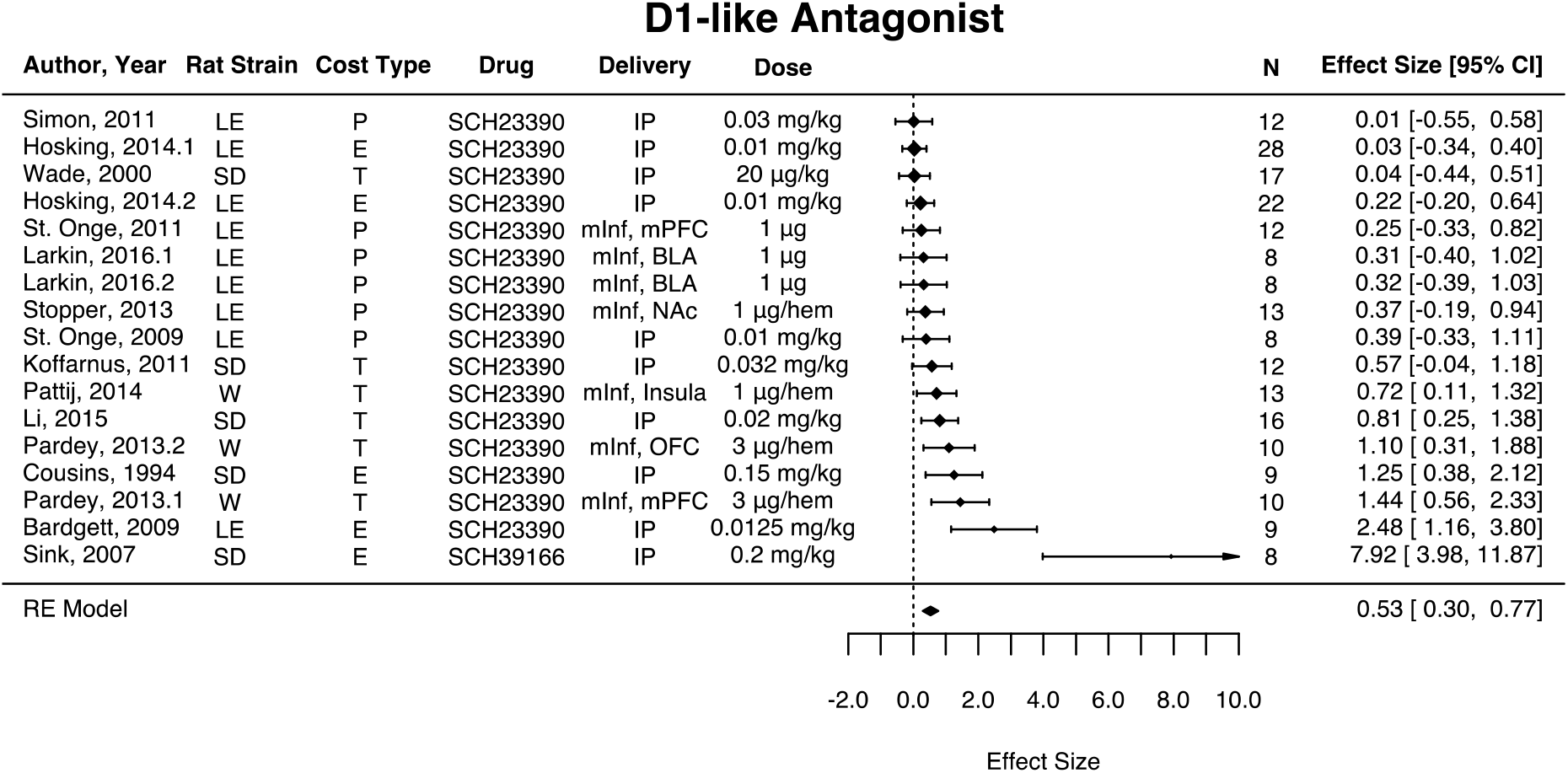
Forest plot of placebo-controlled effect of D1-like antagonism on reward discounting. Positive values indicate increased discounting on drug. LE = Long-Evans, SD = Sprague-Dawley, W = Wistar, P = Probability, T = Time, E = Effort, IP = intraperitoneal, mInf = microinfusion, hem = hemisphere, mPFC = medial prefrontal cortex, NAc = nucleus accumbens, BLA = basolateral amygdala.

#### D2-like agonists

A meta-analysis across D2-like agonists did not identify a significant common effect of drug over placebo on discounting (Q_Heterogeneity_ = 55.1, p < .001, *I^2^* = 74.4%; Cohen’s *d* = .044, SE = .151, 95% CI [-.251, .339], p = .768. Egger’s test for plot asymmetry did not suggest the presence of publication bias (z = −1.67, p = .096). Leave-one-out analysis suggested that removal of any one study had no impact on the overall meta-analytic effect. **See forest plot on Figure 3 and funnel plot on Supplementary Figure S4**.

**Figure 3.**
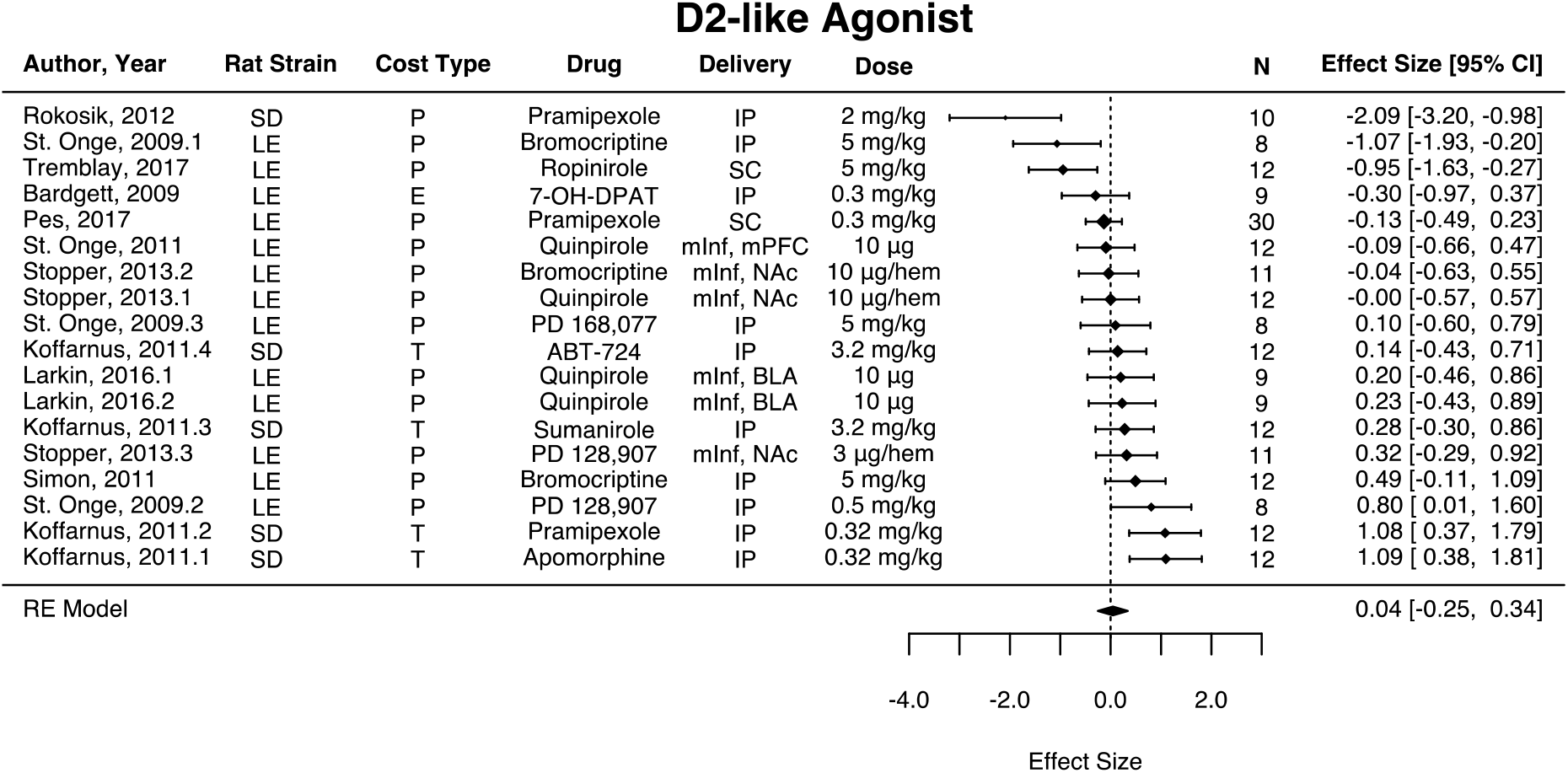
Forest plot of placebo-controlled effect of D2-like agonism on reward discounting. Positive values indicate increased discounting on drug. LE = Long-Evans, SD = Sprague-Dawley, P = Probability, T = Time, E = Effort, IP = intraperitoneal, SC = subcutaneous, mInf = microinfusion, hem = hemisphere, mPFC = medial prefrontal cortex, NAc = nucleus accumbens, BLA = basolateral amygdala.

#### D2-like antagonists

A meta-analysis across D2-like antagonists identified a significant common effect of drug over placebo on discounting (Q_Heterogeneity_ = 166.3, p < .001, *I^2^* = 76.2%; Cohen’s *d* = .505, SE = .097, 95% CI [.315, .696], p < .001. Across effects, D2-like antagonists increased discounting over placebo. Egger’s test for plot asymmetry did suggest the presence of publication bias (z = 8.45, p < .001). Leave-one-out analysis suggested that removal of any one study had no impact on the overall meta-analytic effect. **See forest plot on Figure 4 and funnel plot on Supplementary Figure S5.** Although we included flupenthixol (a drug that antagonizes both D1Rs and D2Rs) effects with D2-like antagonists in the results reported above, excluding flupenthixol effects slightly increases but does not meaningfully alter the effects of D2-like antagonists on discounting (Q_Heterogeneity_ = 129.5, p < .001, *I*^2^ = 84.4%; Cohen’s *d* = .614, SE = .154, 95% CI [.312, .917], p < .001).

**Figure 4.**
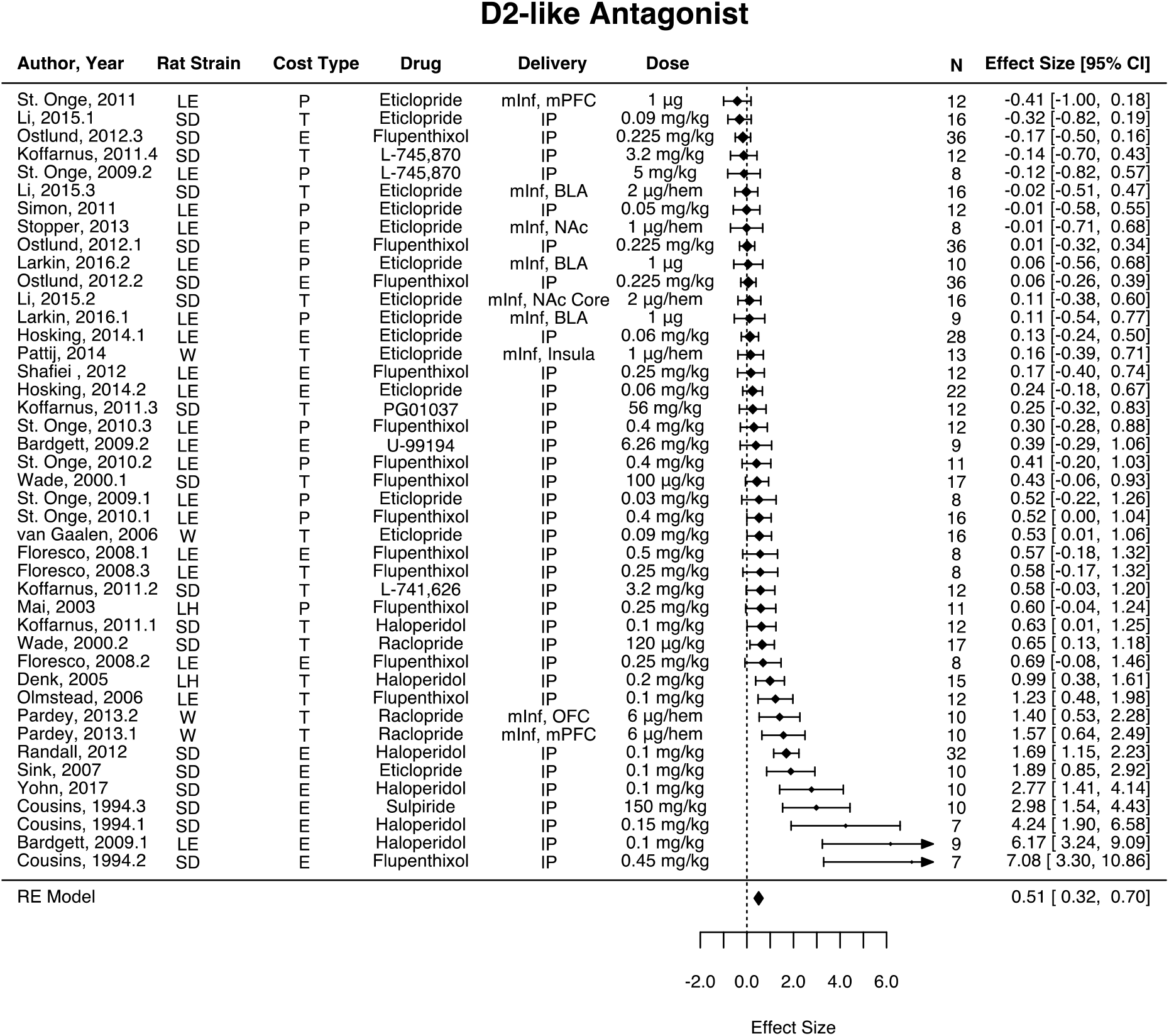
Forest plot of placebo-controlled effect of D2-like antagonism on reward discounting. Positive values indicate increased discounting on drug. LE = Long-Evans, LH = Lister Hooded, SD = Sprague-Dawley, W = Wistar, P = Probability, T = Time, E = Effort, IP = intraperitoneal, mInf = microinfusion, hem = hemisphere, mPFC = medial prefrontal cortex, OFC = orbitofrontal cortex, NAc = nucleus accumbens, BLA = basolateral amygdala.

#### DA transporters

A meta-analysis across DAT-modulating drugs identified a significant common effect of drug over placebo on discounting (Q_Heterogeneity_ = 163.1, p < .001, *I*^2^ = 87.2%; Cohen’s *d* = −.340, SE = .159, 95% CI [−.651, −.028], p = .032. Across effects, DAT-modulating drugs decreased discounting over placebo. Egger’s test for plot asymmetry did not suggest the presence of publication bias (z = −.414, p = .679). Leave-one-out analysis suggested that removal of 9 of the 32 studies impacted the overall meta-analytic effect. The only clear feature that these studies shared were small effect sizes and relatively large variances. As a result, the overall DAT effect on discounting should be interpreted with high caution.

Whereas some drugs like methylphenidate, amphetamine, bupropion, and cocaine have been reported to have substantially higher (and sometimes similar) affinity for DAT over NET (Gatley et al. 1996; Howell and Kimmel 2008; Carroll et al. 2010; Shalabi et al. 2017), atomoxetine and methamphetamine notably have higher affinity for NET than DAT (Rothman et al. 2000, 2001; Bymaster 2002; Howell and Kimmel 2008; Upadhyaya et al. 2013). We therefore repeated the analysis excluding these NET-preferring drugs. Exclusion of effect sizes associated with these drugs (k = 4) indicated, again, that DAT-modulating drugs decreased discounting relative to placebo (Q_Heterogeneity_ = 147.1, p < .001, *I*^2^ = 85.3%; Cohen’s *d* = −.411, SE = .151, 95% CI [−.707, −.115], p = .006). Egger’s test for plot asymmetry did not suggest the presence of publication bias (z = −1.54, p = .125). Here, leave-one-out analysis suggested that removal of any one study had no impact on the overall meta-analytic effect. **See forest plot on Figure 5 and funnel plot on Supplementary Figure S6.**

**Figure 5.**
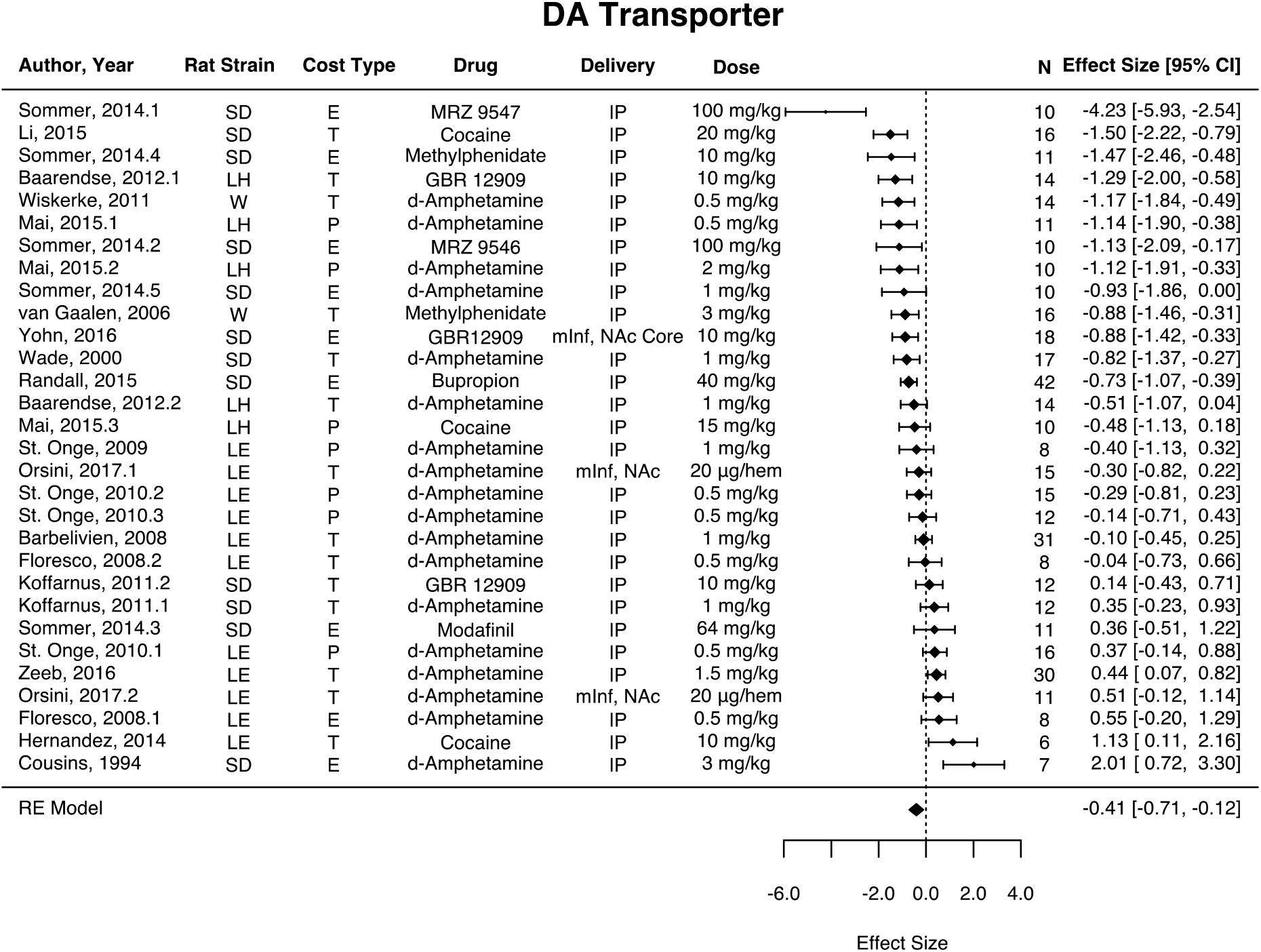
Forest plot of placebo-controlled effect of DAT modulation on reward discounting. Positive values indicate increased discounting on drug. Forest plot excludes effects from NET-preferring drugs. LE = Long-Evans, LH = Lister Hooded, SD = Sprague-Dawley, W = Wistar, P = Probability, T = Time, E = Effort, IP = intraperitoneal, mInf = microinfusion, hem = hemisphere, NAc = nucleus accumbens, BLA = basolateral amygdala.

### Random-effects meta-analysis – exploratory analyses

#### Cost type

A model that included the interaction between reward cost type and drug type suggested a significant moderation effect (Q_Moderator_ = 56.0, p < .001, *I*^2^ = 77.6%). Inspection of the coefficients revealed a main effect within studies of effort discounting (Cohen’s *d* = .915, SE = .338, *p* = .007, 95% CI [.252, 1.58]) and time discounting (Cohen’s *d* = .742, SE = .252, *p* = .003, 95% CI [.248, 1.24]), but not probability discounting (Cohen’s *d* = .153, SE = .190, *p* = .423, 95% CI [−.220, .526]), and a main effect of DAT-modulating drugs (Cohen’s *d* = −1.30, SE = .413, *p* = .002, 95% CI [−2.11, −.488]). There were no significant interactions between reward cost type and drug type.

#### Rat strain

A model that included the interaction between rat strain group and drug type suggested a significant moderation effect (Q_Moderator_ = 71.6, p < .001, *I*^2^ = 71.96%). Inspection of the coefficients revealed a main effect of DAT-modulating drugs *(Cohen’s d* = −3.76, SE = .552, p < .001, 95% CI [−3.15, −.988]) and a significant interaction between DAT-modulating drugs and Long Evans rat strain *(Cohen’s d* = 1.97, SE = .599, p < .01, 95% CI [.793, 3.14]). A follow-up model that tested the effect of rat strain within DAT-modulating drugs suggested a significant effect (Q = 14. 3, p < .007, *I*^2^ = 83.1%) and revealed that discounting for Lister Hooded (Cohen’s *d* = −.896, SE = .352, *p* = .011, 95% CI [−1.59, −.207]) and Sprague-Dawley (Cohen’s *d* = −.436, SE = .220, *p* = .048, 95% CI [−.868, −.004]) rats was significantly reduced on drug over placebo. DAT effects for Wistar (Cohen’s *d* = −1.02, SE = .546, *p* = .061, 95% CI [-2.09, .049]) and Long Evans (Cohen’s *d* = .134, SE = .232, *p* = .562, 95% CI [−.320, −.588]) strains were not statistically significant. **See forest plot on Supplementary Figure S7.**

#### Drug infusion location

From effects that directly infused a DA drug in the brain (k = 28), a model that included the interaction between drug type and infusion location (coded as extra-striatal which includes frontal cortex and amygdala nuclei or striatal which includes infusions within the nucleus accumbens) suggested a weak significant moderation effect (Q_Moderator_ = 11.3, p = .046, *I*^2^ = 38.7%). Inspection of the coefficients revealed a main effect for studies with extra-striatal infusions (Cohen’s *d* = .398, SE = .141, *p* = .005, 95% CI [.121, .674]) but not for striatal infusions (Cohen’s *d* = .229, SE = .275, *p* = .404, 95% CI [−.309, .767]). There were no significant interactions between infusion location and drug type. **See forest plot on Supplementary Figure S8.**

#### Dose-dependent effects

For drugs commonly prescribed to treat Parkinson’s Disease, the levodopa equivalent dose (LED) was estimated for effect sizes associated with D2-like agonists (k = 6) and presynaptic DA agonists (k = 1). A model testing the effect of the continuous LED covariate suggested a significant moderation effect (Q_Moderator_ = 11.54, p < .001, *I^2^* = 75.6%). There was a significant negative correlation between LED and reported effect-size (effect size = −.016, SE = .005, *p* < .001, 95% CI [−.025, −.007]). This means that every 1 mg/kg increase in LED corresponds to a .016 decrease in effect size. **See meta-regression correlation plot on Supplementary Figure S9.** For drugs commonly prescribed to treat psychotic symptoms typically present in schizophrenia and bipolar disorder, the chlorpromazine equivalent dose (CPZ) was estimated for effect sizes associated with D2-like antagonists (k = 21). A model testing the effect of the continuous CPZ covariate did not suggest a significant moderation effect (Q_Moderator_ = 1.08, p = .299, *I*^2^ = 89.9%). There was a non-significant positive correlation between CPZ and reported effect-size (effect size = .017, SE = .017, *p* = .299, 95% CI [-.015, .050]). **See meta-regression correlation plot on Supplementary Figure S10.**

#### D2:D3 receptor affinity

For D2-like agonists and antagonists, the ratio of D2:D3 preferential affinity (determined from prior published reports) was estimated for inclusion as a covariate in meta-regression analyses. Since affinity values were inhibitory constant values (in units of Ki nM), ratios smaller than 1 indicate greater affinity for D2 receptors and ratios larger than 1 indicate greater affinity for D3 receptors. A model for D2-like agonists (k = 13) testing the effect of the continuous D2:D3 affinity ratio covariate did not suggest a significant moderation effect (Q_Moderator_ = .143, p = .705, *I*^2^ = 83.6%). There was no relationship between D2:D3 affinity ratio and reported effect-size (effect size = .003, SE = .008, *p* = .705, 95% CI [-.012, .018]). **See meta-regression correlation plot on Supplementary Figure S11.** A model for D2-like antagonists (k = 24) testing the effect of the continuous D2:D3 affinity ratio covariate did suggest a significant moderation effect (Q_Moderator_ = 25.3, p < .001, *I*^2^ = 70.5%). There was a significant negative relationship between D2:D3 affinity ratio and reported effect-size (Effect size = −3.71, SE = .737,*p* < .001, 95% CI [−5.15, −2.26]). This means that every 1 unit increase in D2:D3 affinity ratio corresponds to a 3.71 decrease in effect size. **See meta-regression correlation plot on Supplementary Figure S12.**

## Discussion

Across 117 effects in rats, a confirmatory quantitative meta-analysis suggested that: (1.) DAT-modulating drugs decrease discounting, (2.) D1-like and D2-like agonists do not impact discounting, and (3.) D1-like and D2-like antagonist moderately increase discounting.

### D1-like and D2-like receptors

The D1-like and D2-like receptor-mediated effects are consistent with emerging perspectives about the relationship between DA receptors and decision making. Traditionally it has been presumed that activation of D1-like and D2-like receptors produces opposing effects (upregulation versus downregulation of intracellular signaling to increase cAMP levels, respectively) in order to support different behaviors (Beaulieu and Gainetdinov 2011). These results could suggest, however, that DA may influence discounting behavior via these receptors in a similar manner. This may be partially consistent with evidence indicating that that D1-like and D2-like receptors engage dissociable processes in the dorsal striatum but not the ventral striatum (Smith et al. 2013; Kupchik et al. 2015; Soares-Cunha et al. 2016; Cox and Witten 2019). A number of convergent findings indicate these receptors are not functionally dissociable in the accumbens for specific motivated behaviors in the ventral striatopallidal pathway (Kupchik et al. 2015; Natsubori et al. 2017; Soares-Cunha et al. 2018). It is therefore possible that reward valuation mechanisms that support discounting behavior are reflected in ventral striatal signaling where activation of D1-like or D2-like receptor types play similar roles. Additional evidence for this D1/D2 similarity comes from our observation that inclusion or exclusion of the D1/D2 antagonist flupenthixol with either D1-like or D2-like receptor antagonists did not substantially change any results. On balance, the meta-analytic effects for D1-like and D2-like receptors do not support a view of opposing direct/indirect pathway function in mediating discounting behavior and suggest that DA receptor antagonism has a greater effect on discounting than agonism.

### DA transporters

The meta-analysis showed that modulation of dopamine transporters moderately decreased discounting behavior. Specifically, increases in DAT-mediated DA release are associated with greater patience, risk aversion, and willingness to expend effort for rewards. These effects are consistent with observations that acute increases in dopamine availability increases motivational vigor for rewards (Westbrook et al. 2020). Although this effect may appear consistent with therapeutic management of ADHD symptoms from DAT-modulating drugs (Volkow et al. 2001, 2011, 2012; Martinez et al. 2020), work from our group and others suggests that we cannot generalize associations between measures of dopamine and discounting across healthy and clinical groups (Castrellon et al. 2019) due to individual differences in baseline DA synthesis capacity and trait impulsivity (Clatworthy et al. 2009; Cools and D’Esposito 2011).

### Reward cost type

Lesion and pharmacological studies in rats have shown that mesolimbic DA similarly impacts probability (St. Onge et al. 2010) and effort (Bardgett et al. 2009) discounting. Complicating this, though, one study has shown that physical and not cognitive effort discounting is modulated by pharmacological stimulation of mesolimbic DA (Hosking et al. 2015). Accordingly, while discounting may exhibit some domain general value processing across cost types, there may be subtle differences in how DA function uniquely accounts for effort requirements in reward preferences. Although it has been assumed that probability and time are discounted similarly (Rachlin et al. 1991), there is evidence that increasing reward magnitude contributes to decreased discounting over time but increased discounting over probabilities (Green et al. 1999). Prior work in humans suggests that the relationship between dopamine function and discounting may vary by cost (time delay, probability, or effort) (Castrellon et al. 2019). Although there was no evidence for differential drug effects across different cost domains in the present meta-analysis, these analyses may have been underpowered. The null effects here do not add substantial evidence to the question of overlap or dissociation in discounting different cost types. It is also critical to recognize that differences in task structure may contribute to differences in behavior. For example, while some studies increased delays with successive trials (ascending procedure), others decreased delays with successive trials (descending procedure) or interchanged delays with successive trials. Although we are underpowered to reliably estimate differences between such effects for each DA site and action, the studies included in our metaanalyses showed some suggestive mean differences in behavior between task procedure types.

There was some evidence for higher mean levels of discounting regardless of drug when an ascending procedure was used compared to a descending or mixed procedure (Barbelivien et al. 2008; St Onge and Floresco 2009; St. Onge et al. 2010; Simon et al. 2011; Shafiei et al. 2012; Randall et al. 2012; Ostlund et al. 2012; Stopper et al. 2013; Larkin et al. 2016; Orsini et al. 2017). Critically, this partial evidence should be interpreted with caution since individual effects varied widely. Further, while some studies reported the time animals spent learning the discounting task structure, others did not. These details are crucial since decisions in later trials may not reflect the same neural or cognitive process that supported decisions in earlier trials. This is of particular relevance here since a number of studies did not identify whether drug and placebo administration order were counterbalanced across animals.

### Rat Strain

An exploratory meta-regression identified an interaction between drug binding site and strain for DAT-modulating drugs. Specifically, within DAT-modulating drugs, Wistar, Lister Hooded, and Sprague Dawley, but not Long Evans rats decreased discounting. The order of rat strain effects (from decreased to increased discounting) indicated that Wistar > Lister Hooded > Sprague Dawley > Long Evans. It has been reported that dopaminergic differences exist between strains (Jiao et al. 2003; Zamudio et al. 2005; Novick et al. 2008; McDermott and Kelly 2008; Rivera-Garcia et al. 2020). Consistent with the meta-analytic effect, Wistar rats have been shown to exhibit higher levels of DAT than Sprague Dawley rats (Zamudio et al. 2005). In addition, inter-strain differences in traits that have been known to covary with dopamine function and motivation like body fat distribution may account for the observed effects (Reed et al. 2011).

### Drug infusion location

An exploratory meta-regression showed that regardless of drug site and action, infusion of substances directly into extrastriatal regions like the medial prefrontal cortex, amygdala, or insula had a greater impact on increasing discounting than those infused into the ventral striatum. In both humans and rats, lesion studies support the importance of both the vmPFC and ventral striatum in discounting behavior. Rats with lesions to the mOFC and NAcc and human patients with vmPFC/OFC lesions discount monetary and food rewards more steeply (Cardinal et al. 2001; Sellitto et al. 2010; Mar et al. 2011). Future work should evaluate whether appreciable differences exist in striatal versus extrastriatal dopaminergic signaling on discounting behavior.

### Dose-dependent effects

Exploratory meta-regressions tested whether drug dose (chlorpromazine equivalent dose for antipsychotic medications and levodopa equivalent dose for anti-Parkinson medications) moderates the effect on discounting. These analyses revealed that variation in presynaptic rescue drug doses were negatively associated with effect sizes. Specifically, studies using lower levodopa equivalent doses (LED) reported higher discounting on drug while studies using higher LED reported lower discounting on drug. This dose-dependency enhances our understanding of the linearity of dopamine effects. It should be cautioned, however, that these dose equivalencies are based on clinical use in humans. Nevertheless, since only some drugs have known LED or CPZ conversion rates, future work should seek to identify an expanded set of dose equivalencies.

### D2:D3 affinity ratio effects

We grouped dopamine receptor drugs into classical D1-like and D2-like families, but it is possible that receptor subtypes exhibit unique effects on behaviors. An exploratory metaregression showed that whereas D2-like agonists had no effect on discounting regardless of preferential affinity for D2 or D3 receptor subtypes, D2-like antagonist effects on increasing discounting are stronger for drugs that have a higher affinity for D2 over D3 receptor subtypes. This is consistent with evidence from a receptor knockout study in mice indicating that D2 but not D3 receptor signaling mediates reinforcing effectiveness of food rewards (Soto et al. 2016) and is complemented by a pharmacological study indicating that D2 but not D3 receptors are critical for attribution of incentive salience to cues (Fraser et al. 2016). However, this is in contrast to pharmacological findings that D2 antagonism attenuates but that D3 antagonism enhances behavioral effects of cocaine (Manvich et al. 2019). Thus, it is not clear from the present data whether D2 and D3 receptors are completely dissociable with respect to drug-mediated effects on reward-related decision making.

### Caveats, Limitations, and Concluding Remarks

Although the meta-analysis yielded important insight about dopaminergic drug effects on discounting behavior, we caution against extrapolating exact claims about function. There are a number of caveats and limitations. First and foremost, although we have a clear understanding of where dopaminergic binding sites are across the brain, drugs do not naturally bind to specific regions. Dopamine receptor signaling varies between the striatum, midbrain, and cortex (Tritsch and Sabatini 2012; Ford 2014) and most drugs are not completely selective for a single receptor subtype (as discussed above) (Closse et al. 1984; Leysen and Gommeren 1986; Wamsley et al. 1991). In addition, although the meta-analysis can explain choice behavior, it cannot speak to the specific value function that supports the underlying behavior. More specifically, it is unclear whether differential modulation of the dopamine system shifts how much weight an animal places on reward magnitudes or costs. This is important because work in rats and humans suggests that costs and magnitudes are anatomically dissociable (Saddoris et al. 2015; Hauser et al. 2017) and one study has shown that methylphenidate effects on humans’ subjective value of cognitive effort depends on caudate DA synthesis capacity (Westbrook et al. 2020). Future work should therefore evaluate whether dopaminergic drugs are more strongly modulating preferences by altering computations supporting integration of costs, magnitudes, and subjective value. In spite of these task and neuropharmacological features, it is notable that there is some degree of consistency across effects—which reflects the ability of meta-analyses to pool heterogenous studies in order to identify generalizable properties.

Since many dopaminergic drugs exhibit non-negligible binding to serotonin, norepinephrine, and adrenergic receptors and transporters, effects cannot be exclusively attributed to dopamine function (Closse et al. 1984; Leysen and Gommeren 1986). In addition, it is important to acknowledge that acute administration of dopaminergic drugs have been demonstrated to have different effects on neurotransmission from chronic administration. For example, acute haloperidol administration contributes to higher spontaneous firing of dopamine neurons than chronic administration (Bunney and Grace 1978). Whereas acute administration of psychostimulants increase dopamine release, PET studies of humans with psychostimulant addictions have shown that chronic use contributes to reduced dopamine release, transporter availability, and D2 receptor availability (Ashok et al. 2017).

The meta-analysis is further limited by the scarcity of studies using primates. Moreover, the literature search and meta-analysis was limited to studies of discounting in healthy animals. While we made this decision to isolate drug effects from disruptions in behavior due to lesions or psychopathology, it is possible that drugs may impact animals depending on systemic alterations to circuits from diseases such as Parkinson’s Disease or schizophrenia (Castrellon et al. 2019; Amlung et al. 2019). An additional caveat raised by the exploratory meta-regressions suggests that drug doses may partially account for reported effect sizes. Since the meta-analysis data extraction protocol was limited to selection of the highest dose effect when multiple doses were available, the effects cannot adequately account for non-linear effects of drug dose on discounting.

While this meta-analysis was limited to discounting paradigms, future meta-studies could evaluate how dopamine pharmacology effects differ for other reward-related behaviors. For example, many studies have evaluated dopaminergic drug effects on reinforcement and probabilistic reversal learning. It is unknown whether consistent cross-study patterns of results with respect to D1-like and D2-like receptors would emerge given that discounted value representations may reflect the same updated valuation process in reinforcement learning, relying on signaling in the ventral striatum and medial prefrontal cortex (Hare et al. 2008). Since behavioral pharmacology meta-analyses typically include only tens of effect sizes, our analysis of 117 effects represents a critical advancement for the field. In general, we hope this metaanalysis encourages additional meta-analytic work in behavioral pharmacology and provision of publicly available data and adoption of practices that standardize reporting critical details. The present meta-analysis contributes to our understanding of how dopamine signaling mediates preferences for delayed, effortful, or uncertain reward outcomes.

## Acknowledgements

J.J.C. was supported by the National Science Foundation Graduate Research Fellowship (grant no. NSF DGE-1644868). We thank Scott A. Huettel, R. Alison Adcock, and John M. Pearson for helpful comments on an earlier draft of the manuscript.

## Data Availability

Data and code used in analyses can be viewed and downloaded on OSF (https://osf.io/27cqw/).

## Author Contributions

J.J.C. and G.R.S.L. conceived and designed the research. J.J.C., J.M., L.G., and K.H. collected the data. J.J.C. analyzed the data with input from G.R.S.L. J.J.C. and G.R.S.L. wrote the manuscript. All authors approved the final version of the manuscript.

## References

Amlung M, Marsden E, Holshausen K, et al (2019) Delay Discounting as a Transdiagnostic Process in Psychiatric Disorders: A Meta-analysis. JAMA Psychiatry 76:1176. https://doi.org/10.1001/jamapsychiatry.2019.2102

Ashok AH, Mizuno Y, Volkow ND, Howes OD (2017) Association of Stimulant Use With Dopaminergic Alterations in Users of Cocaine, Amphetamine, or Methamphetamine: A Systematic Review and Meta-analysis. JAMA Psychiatry 74:511. https://doi.org/10.1001/jamapsychiatry.2017.0135

Barbelivien A, Billy E, Lazarus C, et al (2008) Rats with different profiles of impulsive choice behavior exhibit differences in responses to caffeine and d-amphetamine and in medial prefrontal cortex 5-HT utilization. Behav Brain Res 187:273–283. https://doi.org/10.1016/j.bbr.2007.09.020

Bardgett ME, Depenbrock M, Downs N, et al (2009) Dopamine modulates effort-based decision making in rats. Behav Neurosci 123:242–251. https://doi.org/10.1037/a0014625

Beaulieu J-M, Gainetdinov RR (2011) The Physiology, Signaling, and Pharmacology of Dopamine Receptors. Pharmacol Rev 63:182–217. https://doi.org/10.1124/pr.110.002642

Becker BJ (1988) Synthesizing standardized mean-change measures. Br J Math Stat Psychol 41:257–278. https://doi.org/10.1111/j.2044-8317.1988.tb00901.x

Borenstein M, Hedges LV, Higgins JP, Rothstein HR (2011) Introduction to meta-analysis. John Wiley & Sons

Bunney BS, Grace AA (1978) Acute and chronic haloperidol treatment: Comparison of effects on nigral dopaminergic cell activity. Life Sci 23:1715–1727. https://doi.org/10.1016/0024-3205(78)90471-X

Bymaster F (2002) Atomoxetine Increases Extracellular Levels of Norepinephrine and Dopamine in Prefrontal Cortex of Rat A Potential Mechanism for Efficacy in Attention Deficit/Hyperactivity Disorder. Neuropsychopharmacology 27:699–711. https://doi.org/10.1016/S0893-133X(02)00346-9

Cardinal R, Pennicott D, Sugathapala C, et al (2001) Impulsive Choice Induced in Rats by Lesions of the Nucleus Accumbens Core. Science 292:2499–2501. https://doi.org/10.1126/science.1060818

Carroll FI, Blough BE, Mascarella SW, et al (2010) Synthesis and Biological Evaluation of Bupropion Analogues as Potential Pharmacotherapies for Smoking Cessation. J Med Chem 53:2204–2214. https://doi.org/10.1021/jm9017465

Castrellon JJ, Seaman KL, Crawford JL, et al (2019) Individual Differences in Dopamine Are Associated with Reward Discounting in Clinical Groups But Not in Healthy Adults. J Neurosci 39:321–332. https://doi.org/10.1523/JNEUROSCI.1984-18.2018

Cho DI, Zheng M, Kim K-M (2010) Current perspectives on the selective regulation of dopamine D2 and D3 receptors. Arch Pharm Res 33:1521–1538. https://doi.org/10.1007/s12272-010-1005-8

Clatworthy PL, Lewis SJG, Brichard L, et al (2009) Dopamine Release in Dissociable Striatal Subregions Predicts the Different Effects of Oral Methylphenidate on Reversal Learning and Spatial Working Memory. J Neurosci 29:4690–4696. https://doi.org/10.1523/JNEUROSCI.3266-08.2009

Closse A, Frick W, Dravid A, et al (1984) Classification of drugs according to receptor binding profiles. Naunyn Schmiedebergs Arch Pharmacol 327:95–101. https://doi.org/10.1007/BF00500901

Cools R, D’Esposito M (2011) Inverted-U–Shaped Dopamine Actions on Human Working Memory and Cognitive Control. Biol Psychiatry 69:e113–e125. https://doi.org/10.1016/j.biopsych.2011.03.028

Cousins MS, Wei W, Salamone JD (1994) Pharmacological characterization of performance on a concurrent lever pressing/feeding choice procedure: effects of dopamine antagonist, cholinomimetic, sedative and stimulant drugs. Psychopharmacology (Berl) 116:529–537. https://doi.org/10.1007/BF02247489

Cox J, Witten IB (2019) Striatal circuits for reward learning and decision-making. Nat Rev Neurosci 20:482–494. https://doi.org/10.1038/s41583-019-0189-2

de Wit H, Flory JD, Acheson A, et al (2007) IQ and nonplanning impulsivity are independently associated with delay discounting in middle-aged adults. Personal Individ Differ 42:111–121. https://doi.org/10.1016/j.paid.2006.06.026

Floresco SB, Tse MTL, Ghods-Sharifi S (2008) Dopaminergic and Glutamatergic Regulation of Effort- and Delay-Based Decision Making. Neuropsychopharmacology 33:1966–1979. https://doi.org/10.1038/sj.npp.1301565

Ford CP (2014) The role of D2-autoreceptors in regulating dopamine neuron activity and transmission. Neuroscience 282:13–22. https://doi.org/10.1016/j.neuroscience.2014.01.025

Fraser KM, Haight JL, Gardner EL, Flagel SB (2016) Examining the role of dopamine D2 and D3 receptors in Pavlovian conditioned approach behaviors. Behav Brain Res 305:87–99. https://doi.org/10.1016/j.bbr.2016.02.022

Gatley SJ, Pan D, Chen R, et al (1996) Affinities of methylphenidate derivatives for dopamine, norepinephrine and serotonin transporters. Life Sci 58:PL231–PL239. https://doi.org/10.1016/0024-3205(96)00052-5

Green L, Myerson J, Ostaszewski P (1999) Amount of reward has opposite effects on the discounting of delayed and probabilistic outcomes. J Exp Psychol Learn Mem Cogn 25:418–427. https://doi.org/10.1037/0278-7393.25.2.418

Hare TA, O’Doherty J, Camerer CF, et al (2008) Dissociating the Role of the Orbitofrontal Cortex and the Striatum in the Computation of Goal Values and Prediction Errors. J Neurosci 28:5623–5630. https://doi.org/10.1523/JNEUROSCI.1309-08.2008

Hauser TU, Eldar E, Dolan RJ (2017) Separate mesocortical and mesolimbic pathways encode effort and reward learning signals. Proc Natl Acad Sci 114:E7395–E7404. https://doi.org/10.1073/pnas.1705643114

Hayden BY (2016) Time discounting and time preference in animals: A critical review. Psychon Bull Rev 23:39–53. https://doi.org/10.3758/s13423-015-0879-3

Hedges LV (1989) An Unbiased Correction for Sampling Error in Validity Generalization Studies. J Appl Psychol 74:469–477. https://doi.org/10.1037/0021-9010.74.3.469

Heilbronner SR (2017) Modeling risky decision-making in nonhuman animals: shared core features. Curr Opin Behav Sci 16:23–29. https://doi.org/10.1016/j.cobeha.2017.03.001

Hosking JG, Floresco SB, Winstanley CA (2015) Dopamine Antagonism Decreases Willingness to Expend Physical, But Not Cognitive, Effort: A Comparison of Two Rodent Cost/Benefit Decision-Making Tasks. Neuropsychopharmacology 40:1005–1015. https://doi.org/10.1038/npp.2014.285

Howell LL, Kimmel HL (2008) Monoamine transporters and psychostimulant addiction. Biochem Pharmacol 75:196–217. https://doi.org/10.1016/j.bcp.2007.08.003

Jiao X, Paré WP, Tejani-Butt S (2003) Strain differences in the distribution of dopamine transporter sites in rat brain. Prog Neuropsychopharmacol Biol Psychiatry 27:913–919. https://doi.org/10.1016/S0278-5846(03)00150-7

Koffarnus MN, Newman AH, Grundt P, et al (2011) Effects of selective dopaminergic compounds on a delay-discounting task: Behav Pharmacol 22:300–311. https://doi.org/10.1097/FBP.0b013e3283473bcb

Kupchik YM, Brown RM, Heinsbroek JA, et al (2015) Coding the direct/indirect pathways by D1 and D2 receptors is not valid for accumbens projections. Nat Neurosci 18:1230–1232. https://doi.org/10.1038/nn.4068

Larkin JD, Jenni NL, Floresco SB (2016) Modulation of risk/reward decision making by dopaminergic transmission within the basolateral amygdala. Psychopharmacology (Berl) 233:121–136. https://doi.org/10.1007/s00213-015-4094-8

Lempert KM, Steinglass JE, Pinto A, et al (2019) Can delay discounting deliver on the promise of RDoC? Psychol Med 49:190–199. https://doi.org/10.1017/S0033291718001770

Leysen JE, Gommeren W (1986) Drug-receptor dissociation time, new tool for drug research: Receptor binding affinity and drug-receptor dissociation profiles of serotonin-S2, Dopamine-D2, histamine-H1 antagonists, and opiates. Drug Dev Res 8:119–131. https://doi.org/10.1002/ddr.430080115

Li Y, Zuo Y, Yu P, et al (2015) Role of basolateral amygdala dopamine D2 receptors in impulsive choice in acute cocaine-treated rats. Behav Brain Res 287:187–195. https://doi.org/10.1016/j.bbr.2015.03.039

Löber S, Hübner H, Tschammer N, Gmeiner P (2011) Recent advances in the search for D3 - and D4-selective drugs: probes, models and candidates. Trends Pharmacol Sci 32:148–157. https://doi.org/10.1016/j.tips.2010.12.003

Manvich DF, Petko AK, Branco RC, et al (2019) Selective D2 and D3 receptor antagonists oppositely modulate cocaine responses in mice via distinct postsynaptic mechanisms in nucleus accumbens. Neuropsychopharmacology 44:1445–1455. https://doi.org/10.1038/s41386-019-0371-2

Mar AC, Walker ALJ, Theobald DE, et al (2011) Dissociable Effects of Lesions to Orbitofrontal Cortex Subregions on Impulsive Choice in the Rat. J Neurosci 31:6398–6404. https://doi.org/10.1523/JNEUROSCI.6620-10.2011

Martinez E, Pasquereau B, Drui G, et al (2020) Ventral striatum supports Methylphenidate therapeutic effects on impulsive choices expressed in temporal discounting task. Sci Rep 10:716. https://doi.org/10.1038/s41598-020-57595-6

McDermott C, Kelly JP (2008) Comparison of the behavioural pharmacology of the Lister-Hooded with 2 commonly utilised albino rat strains. Prog Neuropsychopharmacol Biol Psychiatry 32:1816–1823. https://doi.org/10.1016/j.pnpbp.2008.08.004

Moher D (2009) Preferred Reporting Items for Systematic Reviews and Meta-Analyses: The PRISMA Statement. Ann Intern Med 151:264. https://doi.org/10.7326/0003-4819-151-4-200908180-00135

Morris SB, DeShon RP (2002) Combining effect size estimates in meta-analysis with repeated measures and independent-groups designs. Psychol Methods 7:105–125. https://doi.org/10.1037/1082-989X.7.1.105

Natsubori A, Tsutsui-Kimura I, Nishida H, et al (2017) Ventrolateral Striatal Medium Spiny Neurons Positively Regulate Food-Incentive, Goal-Directed Behavior Independently of D1 and D2 Selectivity. J Neurosci 37:2723–2733. https://doi.org/10.1523/JNEUROSCI.3377-16.2017

Nichols DE (2010) Dopamine Receptor Subtype-Selective Drugs: D1-Like Receptors. In: Neve K (ed) The Dopamine Receptors, 2nd edn. Humana Press, Totowa, NJ, pp 75–100

Novick A, Yaroslavsky I, Tejani-Butt S (2008) Strain differences in the expression of dopamine D1 receptors in Wistar–Kyoto (WKY) and Wistar rats. Life Sci 83:74–78. https://doi.org/10.1016/j.lfs.2008.05.006

Orsini CA, Mitchell MR, Heshmati SC, et al (2017) Effects of nucleus accumbens amphetamine administration on performance in a delay discounting task. Behav Brain Res 321:130–136. https://doi.org/10.1016/j.bbr.2017.01.001

Ostlund SB, Kosheleff AR, Maidment NT (2012) Relative Response Cost Determines the Sensitivity of Instrumental Reward Seeking to Dopamine Receptor Blockade. Neuropsychopharmacology 37:2653–2660. https://doi.org/10.1038/npp.2012.129

Pardey MC, Kumar NN, Goodchild AK, Cornish JL (2013) Catecholamine receptors differentially mediate impulsive choice in the medial prefrontal and orbitofrontal cortex. J Psychopharmacol (Oxf) 27:203–212. https://doi.org/10.1177/0269881112465497

Pattij T, Vanderschuren L (2008) The neuropharmacology of impulsive behaviour. Trends Pharmacol Sci 29:192–199. https://doi.org/10.1016/j.tips.2008.01.002

Rachlin H, Raineri A, Cross D (1991) Subjective Probability and Delay. J Exp Anal Behav 55:233–244. https://doi.org/10.1901/jeab.1991.55-233

Randall PA, Pardo M, Nunes EJ, et al (2012) Dopaminergic Modulation of Effort-Related Choice Behavior as Assessed by a Progressive Ratio Chow Feeding Choice Task: Pharmacological Studies and the Role of Individual Differences. PLoS ONE 7:e47934. https://doi.org/10.1371/journal.pone.0047934

Reed DR, Duke FF, Ellis HK, et al (2011) Body fat distribution and organ weights of 14 common strains and a 22-strain consomic panel of rats. Physiol Behav 103:523–529. https://doi.org/10.1016/j.physbeh.2011.04.006

Reimers S, Maylor EA, Stewart N, Chater N (2009) Associations between a one-shot delay discounting measure and age, income, education and real-world impulsive behavior. Personal Individ Differ 47:973–978. https://doi.org/10.1016/j.paid.2009.07.026

Rivera-Garcia MT, McCane AM, Chowdhury TG, et al (2020) Sex and strain differences in dynamic and static properties of the mesolimbic dopamine system. Neuropsychopharmacology. https://doi.org/10.1038/s41386-020-0765-1

Rohatgi, Ankit (2019) WebPlotDigitizer. Version 4.2. San Francisco, California, USA. URL https://automeris.io/WebPlotDigitizer

Rosati AG, Hare B (2016) Reward currency modulates human risk preferences. Evol Hum Behav 37:159–168. https://doi.org/10.1016/j.evolhumbehav.2015.10.003

Rothman RB, Baumann MH, Dersch CM, et al (2001) Amphetamine-type central nervous system stimulants release norepinephrine more potently than they release dopamine and serotonin. Synapse 39:32–41. https://doi.org/10.1002/1098-2396(20010101)39:1<32::AID-SYN5>3.0.CO;2-3

Rothman RB, Partilla JS, Baumann MH, et al (2000) Neurochemical neutralization of methamphetamine with high-affinity nonselective inhibitors of biogenic amine transporters: a pharmacological strategy for treating stimulant abuse. Synapse 35:222–227. https://doi.org/10.1002/(SICI)1098-2396(20000301)35:3<222::AID-SYN7>3.0.CO;2-K

Saddoris MP, Sugam JA, Stuber GD, et al (2015) Mesolimbic Dopamine Dynamically Tracks, and Is Causally Linked to, Discrete Aspects of Value-Based Decision Making. Biol Psychiatry 77:903–911. https://doi.org/10.1016/j.biopsych.2014.10.024

Seaman KL, Gorlick MA, Vekaria KM, et al (2016) Adult age differences in decision making across domains: Increased discounting of social and health-related rewards. Psychol Aging 31:737–746. https://doi.org/10.1037/pag0000131

Sellitto M, Ciaramelli E, di Pellegrino G (2010) Myopic Discounting of Future Rewards after Medial Orbitofrontal Damage in Humans. J Neurosci 30:16429–16436. https://doi.org/10.1523/JNEUROSCI.2516-10.2010

Shafiei N, Gray M, Viau V, Floresco SB (2012) Acute Stress Induces Selective Alterations in Cost/Benefit Decision-Making. Neuropsychopharmacology 37:2194–2209. https://doi.org/10.1038/npp.2012.69

Shalabi AR, Walther D, Baumann MH, Glennon RA (2017) Deconstructed Analogues of Bupropion Reveal Structural Requirements for Transporter Inhibition versus Substrate-Induced Neurotransmitter Release. ACS Chem Neurosci 8:1397–1403. https://doi.org/10.1021/acschemneuro.7b00055

Simon NW, Montgomery KS, Beas BS, et al (2011) Dopaminergic Modulation of Risky Decision-Making. J Neurosci 31:17460–17470. https://doi.org/10.1523/JNEUROSCI.3772-11.2011

Smith RJ, Lobo MK, Spencer S, Kalivas PW (2013) Cocaine-induced adaptations in D1 and D2 accumbens projection neurons (a dichotomy not necessarily synonymous with direct and indirect pathways). Curr Opin Neurobiol 23:546–552. https://doi.org/10.1016/j.conb.2013.01.026

Soares-Cunha C, Coimbra B, Domingues AV, et al (2018) Nucleus Accumbens Microcircuit Underlying D2-MSN-Driven Increase in Motivation. eneuro 5:ENEURO.0386-18.2018. https://doi.org/10.1523/ENEURO.0386-18.2018

Soares-Cunha C, Coimbra B, Sousa N, Rodrigues AJ (2016) Reappraising striatal D1- and D2-neurons in reward and aversion. Neurosci Biobehav Rev 68:370–386. https://doi.org/10.1016/j.neubiorev.2016.05.021

Sommer S, Danysz W, Russ H, et al (2014) The dopamine reuptake inhibitor MRZ-9547 increases progressive ratio responding in rats. Int J Neuropsychopharmacol 17:2045–2056. https://doi.org/10.1017/S1461145714000996

Soto PL, Hiranita T, Xu M, et al (2016) Dopamine D2-Like Receptors and Behavioral Economics of Food Reinforcement. Neuropsychopharmacology 41:971–978. https://doi.org/10.1038/npp.2015.223

St. Onge JR, Chiu YC, Floresco SB (2010) Differential effects of dopaminergic manipulations on risky choice. Psychopharmacology (Berl) 211:209–221. https://doi.org/10.1007/s00213-010-1883-y

St Onge JR, Floresco SB (2009) Dopaminergic Modulation of Risk-Based Decision Making. Neuropsychopharmacology 34:681–697. https://doi.org/10.1038/npp.2008.121

Stopper CM, Khayambashi S, Floresco SB (2013) Receptor-Specific Modulation of Risk-Based Decision Making by Nucleus Accumbens Dopamine. Neuropsychopharmacology 38:715–728. https://doi.org/10.1038/npp.2012.240

Susini I, Safryghin A, Hillemann F, Wascher CAF (2020) Delay of gratification in non-human animals: A review of inter- and intra-specific variation in performance. Animal Behavior and Cognition

Tritsch NX, Sabatini BL (2012) Dopaminergic Modulation of Synaptic Transmission in Cortex and Striatum. Neuron 76:33–50. https://doi.org/10.1016/j.neuron.2012.09.023

Upadhyaya HP, Desaiah D, Schuh KJ, et al (2013) A review of the abuse potential assessment of atomoxetine: a nonstimulant medication for attention-deficit/hyperactivity disorder. Psychopharmacology (Berl) 226:189–200. https://doi.org/10.1007/s00213-013-2986-z

Viechtbauer W (2010) Conducting meta-analyses in R with the metafor package. J Stat Softw 36:1–48

Volkow ND, Wang G-J, Fowler JS, et al (2001) Therapeutic Doses of Oral Methylphenidate Significantly Increase Extracellular Dopamine in the Human Brain. J Neurosci 21:RC121–RC121. https://doi.org/10.1523/JNEUROSCI.21-02-j0001.2001

Volkow ND, Wang G-J, Newcorn JH, et al (2011) Motivation deficit in ADHD is associated with dysfunction of the dopamine reward pathway. Mol Psychiatry 16:1147–1154. https://doi.org/10.1038/mp.2010.97

Volkow ND, Wang G-J, Tomasi D, et al (2012) Methylphenidate-Elicited Dopamine Increases in Ventral Striatum Are Associated with Long-Term Symptom Improvement in Adults with Attention Deficit Hyperactivity Disorder. J Neurosci 32:841–849. https://doi.org/10.1523/JNEUROSCI.4461-11.2012

Wade TR, de Wit H, Richards JB (2000) Effects of dopaminergic drugs on delayed reward as a measure of impulsive behavior in rats. Psychopharmacology (Berl) 150:90–101. https://doi.org/10.1007/s002130000402

Wamsley JK, Hunt ME, McQuade RD, Alburges ME (1991) [3H]SCH39166, a D1 dopamine receptor antagonist: Binding characteristics and localization. Exp Neurol 111:145–151. https://doi.org/10.1016/0014-4886(91)90001-S

Westbrook A, van den Bosch R, Määttä JI, et al (2020) Dopamine promotes cognitive effort by biasing the benefits versus costs of cognitive work. Science 367:1362–1366. https://doi.org/10.1126/science.aaz5891

Wetzel H, Gründer G, Hillert A, et al (1998) Amisulpride versus flupentixol in schizophrenia with predominantly positive symptomatology - a double-blind controlled study comparing a selective D 2-like antagonist to a mixed D 1-/D 2-like antagonist. Psychopharmacology (Berl) 137:223–232. https://doi.org/10.1007/s002130050614

Yates JR, Perry JL, Meyer AC, et al (2014) Role of medial prefrontal and orbitofrontal monoamine transporters and receptors in performance in an adjusting delay discounting procedure. Brain Res 1574:26–36. https://doi.org/10.1016/j.brainres.2014.06.004

Zamudio S, Fregoso T, Miranda A, et al (2005) Strain differences of dopamine receptor levels and dopamine related behaviors in rats. Brain Res Bull 65:339–347. https://doi.org/10.1016/j.brainresbull.2005.01.009

