## Supplementary Figure S1 for "Dopaminergic modulation of reward discounting in healthy rats: a systematic review and meta-analysis"

### Supplementary Methods

*Exploratory Meta-analyses*

*Rat strain analysis.* Since the studies evaluated in the meta-analysis included multiple rat strains exposed to drugs that act on different DA binding sites, we ran exploratory meta-analytic models to evaluate whether effect sizes depended on the interaction of these terms. Significant interactions were followed up with models to evaluate the simple main effect of rat strain within DA drug binding site to aide visualization of the effect.

*Discounting cost analysis.* Since the studies evaluated included discounting of time, probability, and effort in response to drugs that bind to different DA binding sites, we ran exploratory meta-analytic models to evaluate whether effect sizes depended on the interaction of these terms. Significant interactions were followed up with models to evaluate the simple main effect of cost type within DA drug binding site to aide visualization of the effect.

*Drug infusion location analysis.* While intraperitoneal administration was the most common drug delivery method across included studies, researchers interested in drug effects on specific brain regions may administer doses to rats via direct infusion. We therefore ran exploratory meta-analyses of these kinds of studies to assess whether drug effects depended on the interaction between DA binding site and infusion location. Locations were coded as extrastriatal or within the nucleus accumbens. The only infusion sites outside the nucleus accumbens [1–4] included the medial prefrontal cortex or orbitofrontal cortex [5–7], basolateral amygdala [3,8], and insula [9]. Significant interactions were followed up with models to evaluate the simple main effect of infusion location within DA drug binding site to aide visualization of the effect.

*Drug dose meta-regressions.* We sought to explore dose-dependent effects of DA drugs on discounting. Specifically, antipsychotic medication and anti-Parkinson's medication have known dose-equivalencies to quantitatively compare different drugs. Using published guidelines based on therapeutic effects in humans [10–14], we calculated the chlorpromazine equivalent dose (CPZ) for antipsychotics and levodopa-equivalent dose (LED) for anti-Parkinson's medication. Meta-regressions were run using either the CPZ or LED as a continuous covariate.

*D2:D3 receptor affinity analysis*. Few drugs are truly selective for a single receptor subtype. In particular, a number of studies have reported that drugs which largely bind to dopamine D2-like receptors show preferential affinity to either D2 or D3 receptor subtypes. These differences are important because not only are D2 and D3 receptors differentially expressed across striatal subregions [15], but their functions have also been differentially associated with a variety of motivated behaviors such as reinforcing effectiveness of food rewards [16]. We therefore explored whether D2-like agonist or antagonist effects on discounting depended on the degree to which the drug preferentially binds to D2 or D3 receptors. To do this, we first searched the literature to identify studies reporting affinity values (inhibitory constants in units of Ki nM) across receptor subtypes for as many drugs we could find. In many cases we identified more than one affinity value for each receptor. We include this supplementary data (with linked references) with all shared data on OSF (https://osf.io/27cqw/). Data include affinity values for D1, D2, D3, D4, and D5 receptors whenever available. Next, for each reported affinity, we calculated a D2:D3 affinity ratio (where lower values indicate greater D2 affinity). Following this, to account for differences in affinities reported in past studies, we first averaged D2:D3 ratios across reports for each drug when more than one ratio was available. We then used this average D2:D3 affinity ratio as a continuous covariate in a meta-regression for D2 agonists and D2 antagonists. For D2 agonists, 2 effect sizes from one drug (PD 128,907) were clear outliers (mean D2:D3 affinity ratio = 626.26). **See histogram plots with and without outliers in Figure S13 below**. We therefore excluded these two outlier effect sizes prior to meta-regression. Similarly, for D2 antagonists, 2 effect sizes related to PG0137 and U-99194 were clear outliers (mean D2:D3 affinity ratios = 133.29 and 10.23, respectively). **See histogram plots with and without outliers in Figure S14 below**. All other antagonist affinity ratios were lower than 1. Including outliers did not change statistical inference for D2 agonists (Effect size = .001, SE = .001, p = .252, 95% CI [-.001, .003]). However D2 antagonist D2:D3 affinity ratios were not related effect sizes when PG0137 and U-99194 effects were included (Effect size = -.004, SE = .007, p = .561, 95% CI [-.017, .009]).

**Figure S1.** PRISMA (Preferred Reporting Items for Systematic Reviews and Meta-Analyses) diagram showing study identification.

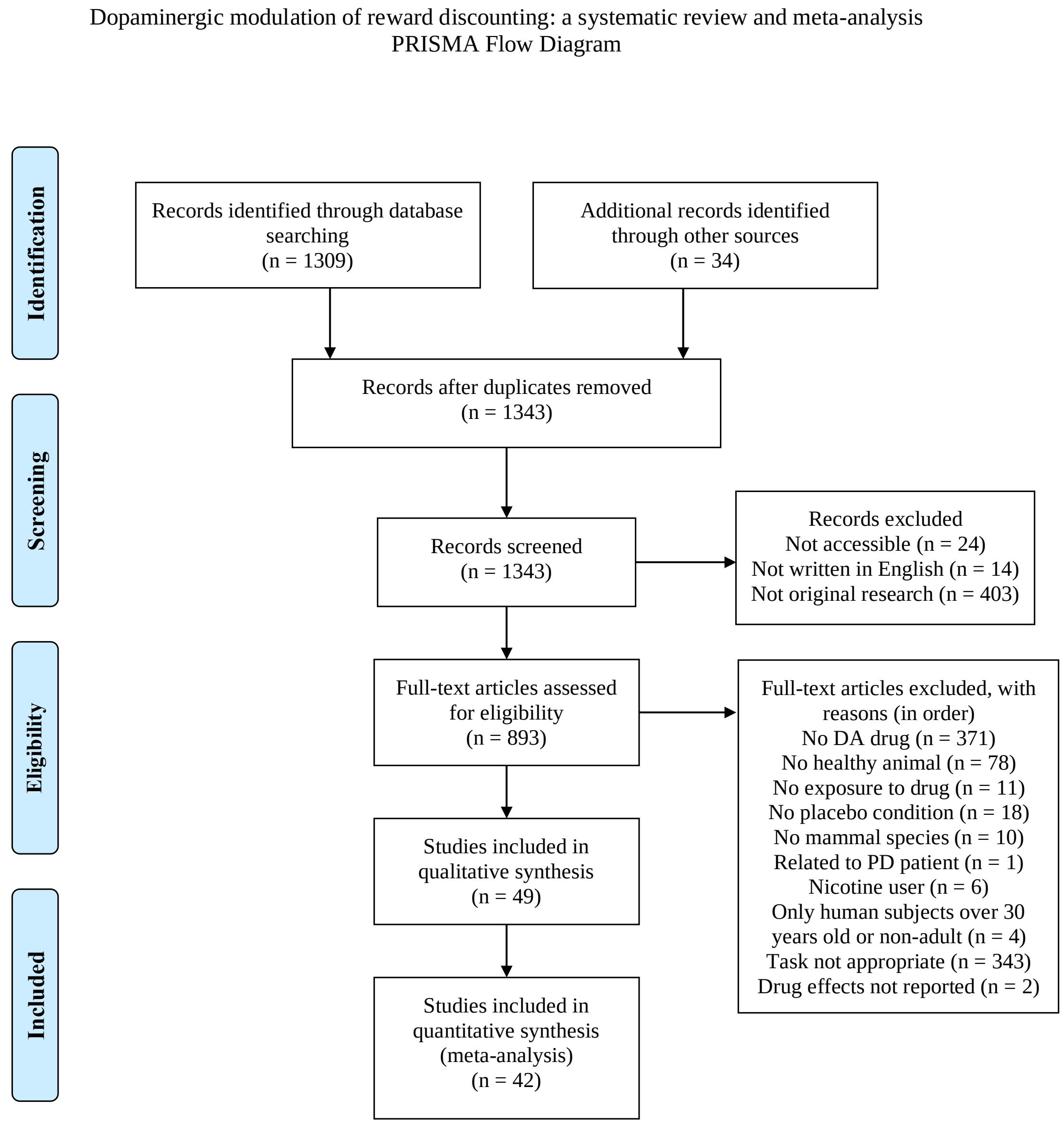

**Figure S2.** Funnel plot of D1-like agonist effect sizes and associated standard errors. The funnel plot is centered at the null-effect value (0). The unshaded region corresponds to p-values greater than .10, the dark gray-shaded region corresponds to p-values between .10 and .05, the medium gray-shaded region corresponds to p-values between .05 and .01, and the region outside of the funnel corresponds to p-values below .01.

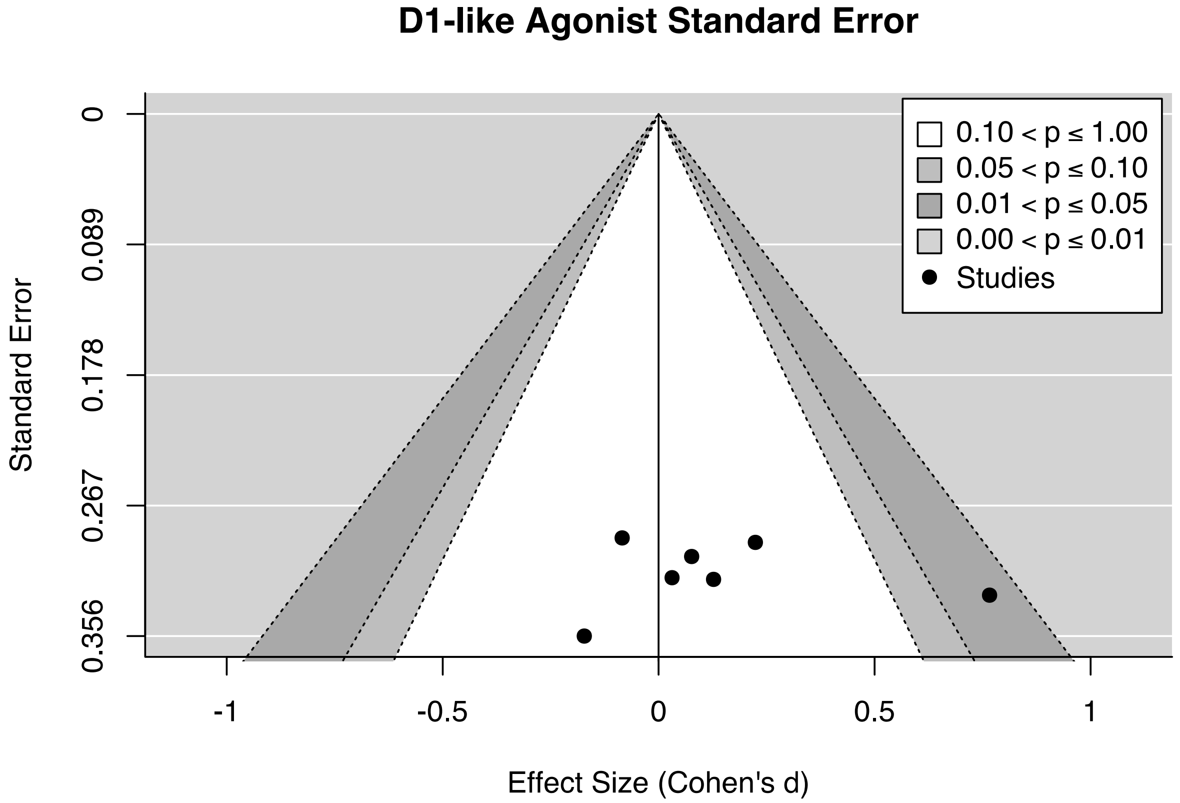

**Figure S3.** Funnel plot of D1-like antagonist effect sizes and associated standard errors. The funnel plot is centered at the null-effect value (0). The unshaded region corresponds to p-values greater than .10, the dark gray-shaded region corresponds to p-values between .10 and .05, the medium gray-shaded region corresponds to p-values between .05 and .01, and the region outside of the funnel corresponds to p-values below .01.

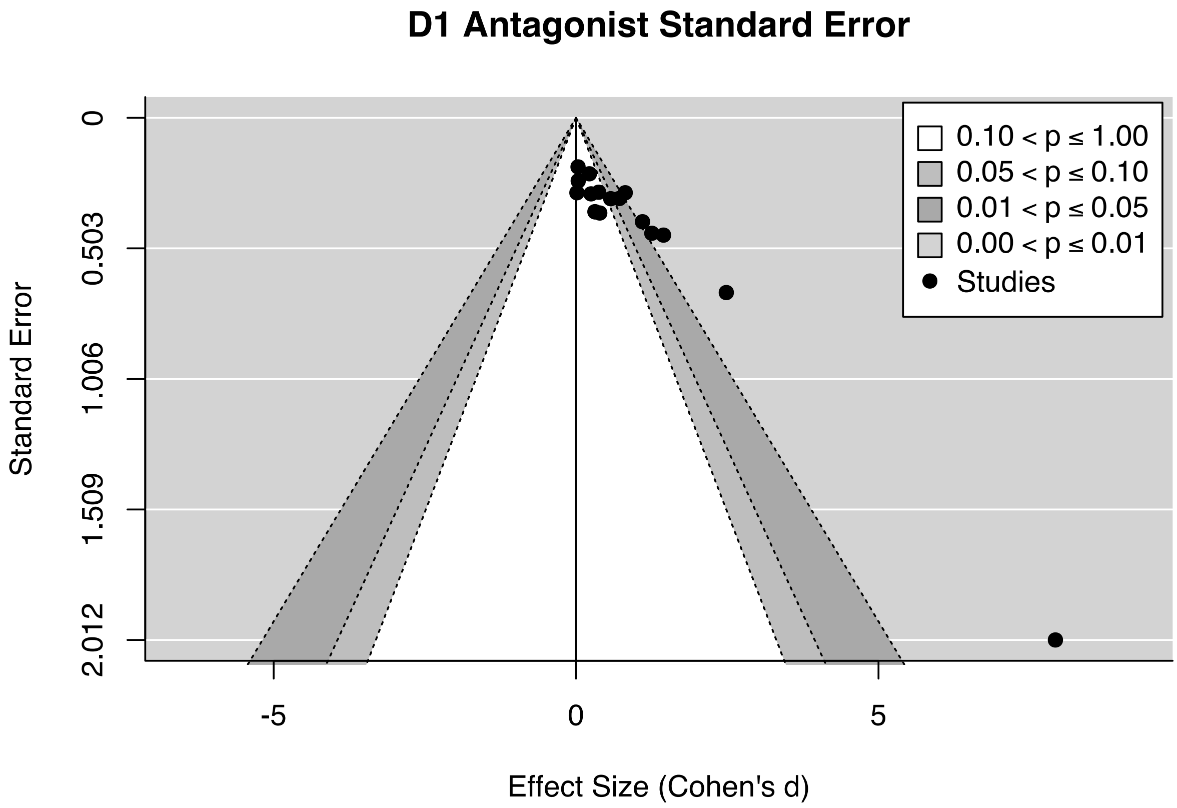

**Figure S4.** Funnel plot of D2-like agonist effect sizes and associated standard errors. The funnel plot is centered at the null-effect value (0). The unshaded region corresponds to p-values greater than .10, the dark gray-shaded region corresponds to p-values between .10 and .05, the medium gray-shaded region corresponds to p-values between .05 and .01, and the region outside of the funnel corresponds to p-values below .01.

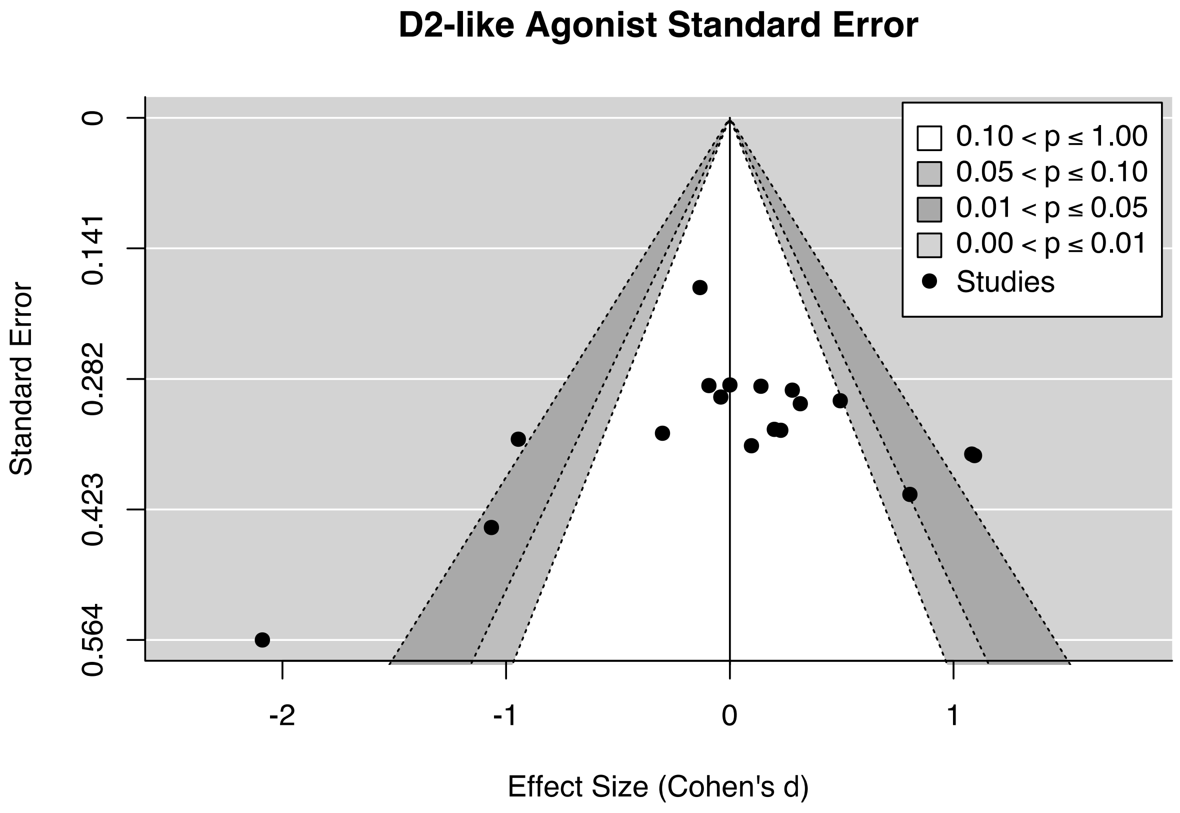

**Figure S5.** Funnel plot of D2-like antagonist effect sizes and associated standard errors. The funnel plot is centered at the null-effect value (0). The unshaded region corresponds to p-values greater than .10, the dark gray-shaded region corresponds to p-values between .10 and .05, the medium gray-shaded region corresponds to p-values between .05 and .01, and the region outside of the funnel corresponds to p-values below .01.

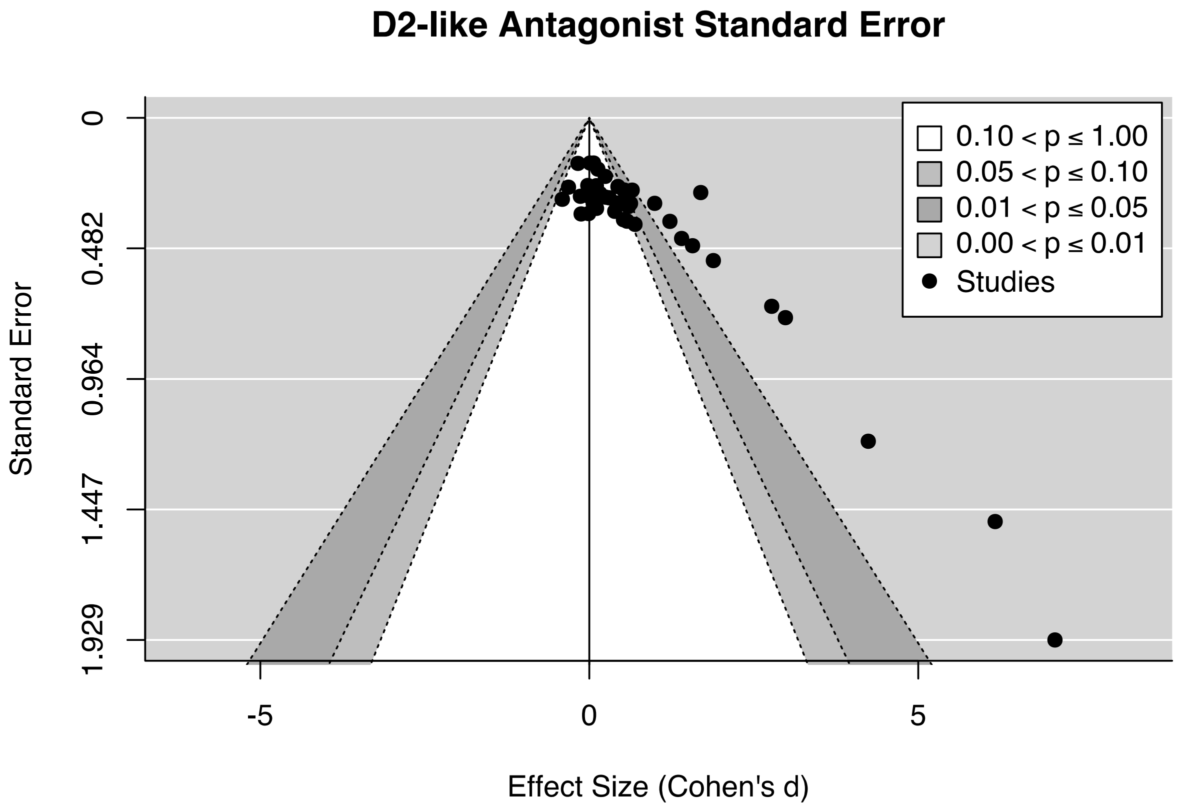

**Figure S6.** Funnel plot of DAT modulation effect sizes and associated standard errors. The funnel plot is centered at the null-effect value (0). The unshaded region corresponds to p-values greater than .10, the dark gray-shaded region corresponds to p-values between .10 and .05, the medium gray-shaded region corresponds to p-values between .05 and .01, and the region outside of the funnel corresponds to p-values below .01. Funnel plot excludes effects from NET-preferring drugs.

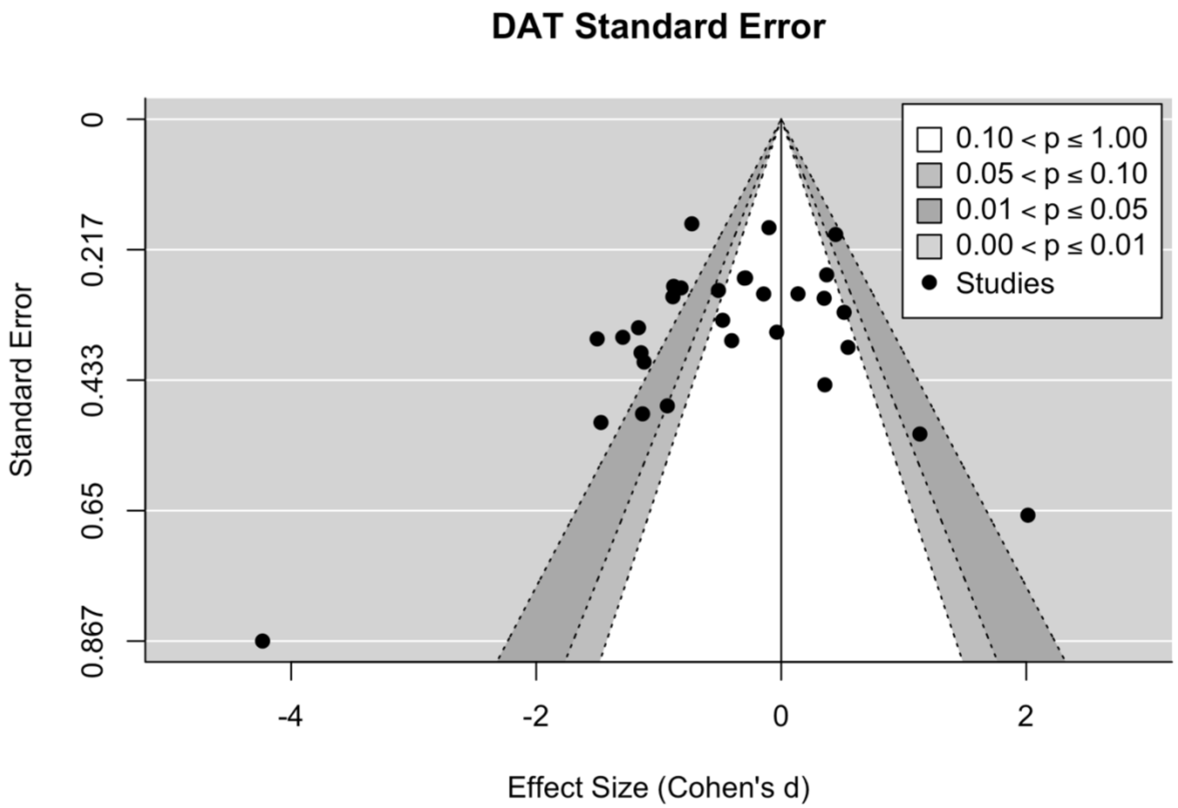

**Figure S7.** Meta-regression forest plot of DAT modulation on reward discounting by rat strain. Shaded polygons indicate the predicted effect for each rat strain. Positive values indicate increased discounting on drug. LE = Long Evans, LH = Lister Hooded, SD = Sprague-Dawley, W = Wistar, P = Probability, T = Time, E = Effort, IP = intraperitoneal, mInf = microinfusion, hem = hemisphere, NAc = nucleus accumbens, BLA = basolateral amygdala.

**DA Transporter**

| Author, Year | Rat Strain | Cost Type | Drug | Delivery | Dose | N | Effect Size [95% CI] |
|---|---|---|---|---|---|---|---|
| St. Onge, 2009 | LE | P | d-Amphetamine | IP | 1 mg/kg | 8 | -0.40 [-1.13, 0.32] |
| Orsini, 2017.1 | LE | T | d-Amphetamine | mInf, NAc | 20 µg/hem | 15 | -0.30 [-0.82, 0.22] |
| St. Onge, 2010.2 | LE | P | d-Amphetamine | IP | 0.5 mg/kg | 15 | -0.29 [-0.81, 0.23] |
| St. Onge, 2010.3 | LE | P | d-Amphetamine | IP | 0.5 mg/kg | 12 | -0.14 [-0.71, 0.43] |
| Barbelivien, 2008 | LE | T | d-Amphetamine | IP | 1 mg/kg | 31 | -0.10 [-0.45, 0.25] |
| Floresco, 2008.2 | LE | T | d-Amphetamine | IP | 0.5 mg/kg | 8 | -0.04 [-0.73, 0.66] |
| St. Onge, 2010.1 | LE | P | d-Amphetamine | IP | 0.5 mg/kg | 16 | 0.37 [-0.14, 0.88] |
| Zeeb, 2016 | LE | T | d-Amphetamine | IP | 1.5 mg/kg | 30 | 0.44 [ 0.07, 0.82] |
| Orsini, 2017.2 | LE | T | d-Amphetamine | mInf, NAc | 20 µg/hem | 11 | 0.51 [-0.12, 1.14] |
| Floresco, 2008.1 | LE | E | d-Amphetamine | IP | 0.5 mg/kg | 8 | 0.55 [-0.20, 1.29] |
| Hernandez, 2014 | LE | T | Cocaine | IP | 10 mg/kg | 6 | 1.13 [ 0.11, 2.16] |
| Baarendse, 2012.1 | LH | T | GBR 12909 | IP | 10 mg/kg | 14 | -1.29 [-2.00, -0.58] |
| Mai, 2015.1 | LH | P | d-Amphetamine | IP | 0.5 mg/kg | 11 | -1.14 [-1.90, -0.38] |
| Mai, 2015.2 | LH | P | d-Amphetamine | IP | 2 mg/kg | 10 | -1.12 [-1.91, -0.33] |
| Baarendse, 2012.2 | LH | T | d-Amphetamine | IP | 1 mg/kg | 14 | -0.51 [-1.07, 0.04] |
| Mai, 2015.3 | LH | P | Cocaine | IP | 15 mg/kg | 10 | -0.48 [-1.13, 0.18] |
| Sommer, 2014.1 | SD | E | MRZ 9547 | IP | 100 mg/kg | 10 | -4.23 [-5.93, -2.54] |
| Li, 2015 | SD | T | Cocaine | IP | 20 mg/kg | 16 | -1.50 [-2.22, -0.79] |
| Sommer, 2014.4 | SD | E | Methylphenidate | IP | 10 mg/kg | 11 | -1.47 [-2.46, -0.48] |
| Sommer, 2014.2 | SD | E | MRZ 9546 | IP | 100 mg/kg | 10 | -1.13 [-2.09, -0.17] |
| Sommer, 2014.5 | SD | E | d-Amphetamine | IP | 1 mg/kg | 10 | -0.93 [-1.86, 0.00] |
| Yohn, 2016.1 | SD | E | GBR12909 | mInf, NAc Core | 10 mg/kg | 18 | -0.88 [-1.42, -0.33] |
| Wade, 2000 | SD | T | d-Amphetamine | IP | 1 mg/kg | 17 | -0.82 [-1.37, -0.27] |
| Randall, 2015 | SD | E | Bupropion | IP | 40 mg/kg | 42 | -0.73 [-1.07, -0.39] |
| Siemian, 2017 | SD | T | d-Methamphetamine | IP | 1 mg/kg | 8 | 0.03 [-0.67, 0.72] |
| Koffarnus, 2011.2 | SD | T | GBR 12909 | IP | 10 mg/kg | 12 | 0.14 [-0.43, 0.71] |
| Koffarnus, 2011.1 | SD | T | d-Amphetamine | IP | 1 mg/kg | 12 | 0.35 [-0.23, 0.93] |
| Sommer, 2014.3 | SD | E | Modafinil | IP | 64 mg/kg | 11 | 0.36 [-0.51, 1.22] |
| Cousins, 1994 | SD | E | d-Amphetamine | IP | 3 mg/kg | 7 | 2.01 [ 0.72, 3.30] |
| Yohn, 2016.2 | SD | E | Atomoxetine | IP | 1 mg/kg | 8 | 2.31 [ 0.98, 3.64] |
| Wiskerke, 2011 | W | T | d-Amphetamine | IP | 0.5 mg/kg | 14 | -1.17 [-1.84, -0.49] |
| van Gaalen, 2006 | W | T | Methylphenidate | IP | 3 mg/kg | 16 | -0.88 [-1.46, -0.31] |

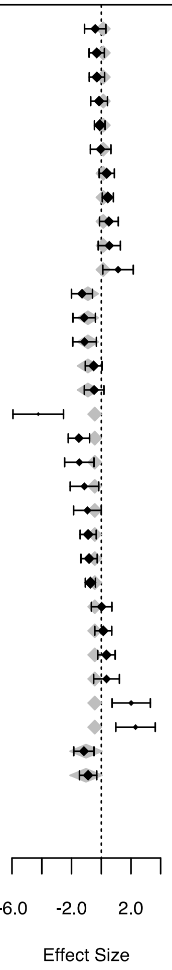

**Figure S8.** Meta-regression forest plot of placebo-controlled effect of drug infusion location on reward discounting. Shaded polygons indicate the predicted effect for each infusion location. Positive values indicate increased discounting on drug. LE = Long Evans, LH = Lister Hooded, SD = Sprague-Dawley, W = Wistar, P = Probability, T = Time, E = Effort, IP = intraperitoneal, hem = hemisphere.

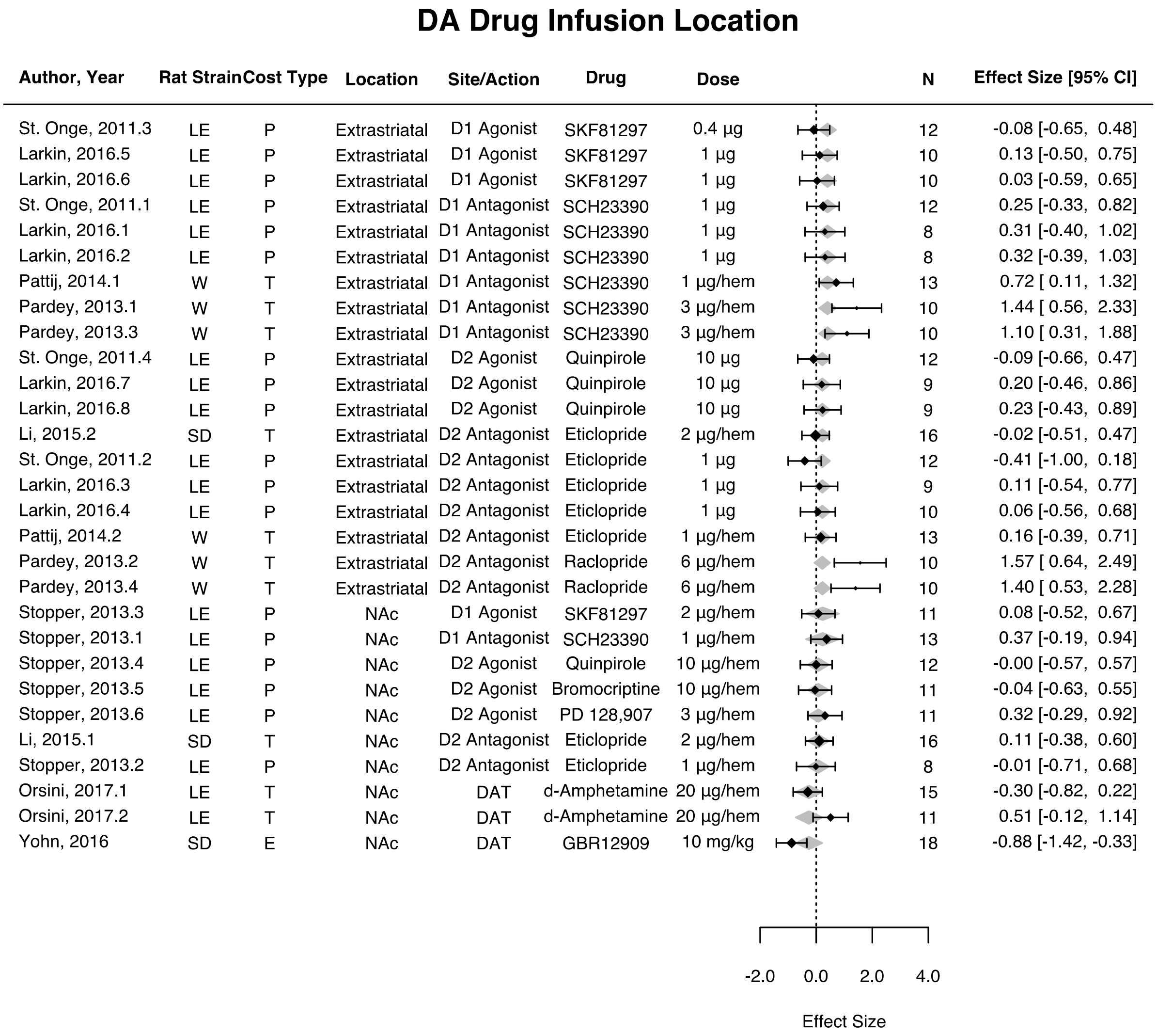

**Figure S9.** Meta-regression of placebo-controlled effect of anti-Parkinson levodopa equivalent dose (LED) on reward discounting showing predicted regression slope and 95% confidence interval. Points represent individual effect sizes scaled by the ratio of the effect size to standard error. Positive effect sizes indicate increased discounting on drug.

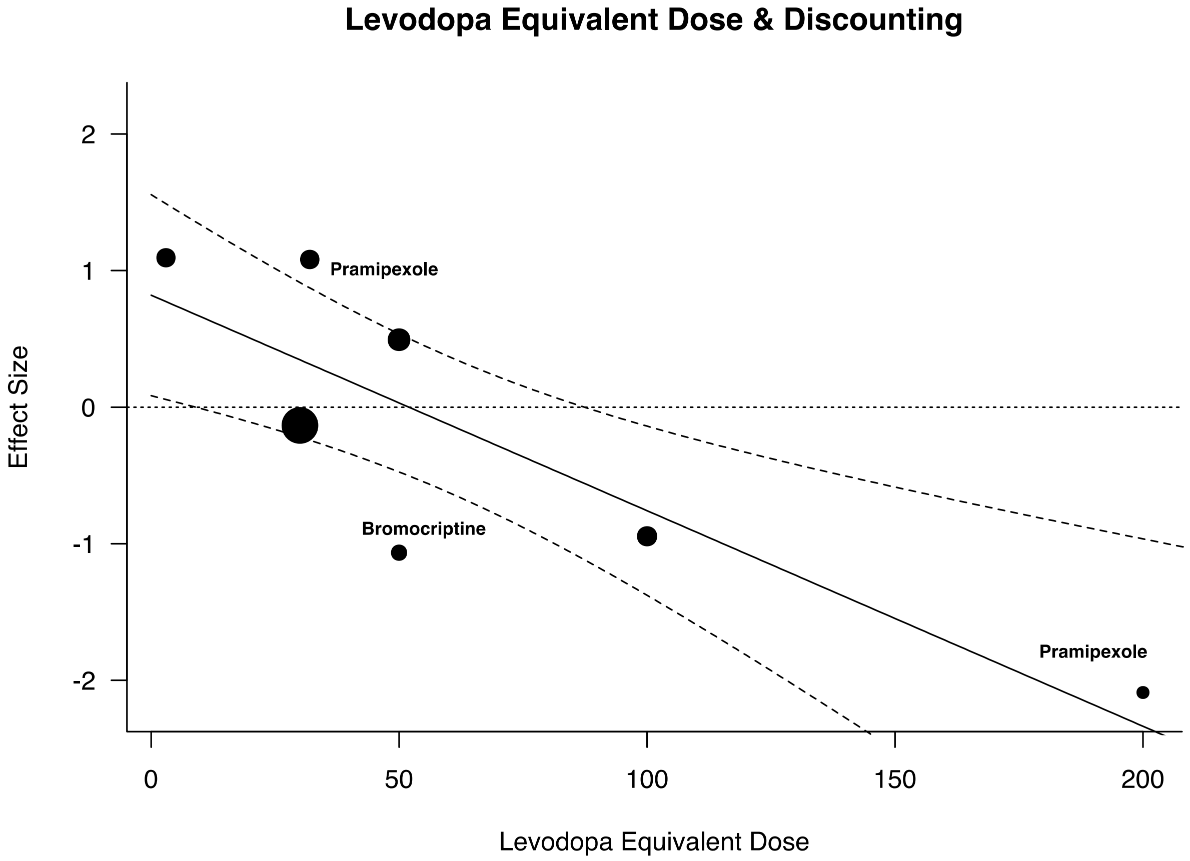

**Figure S10.** Meta-regression of placebo-controlled effect of antipsychotic chlorpromazine equivalent dose (CPZ) on reward discounting showing predicted regression slope and 95% confidence interval. Points represent individual effect sizes scaled by the ratio of the effect size to standard error. Positive effect sizes indicate increased discounting on drug.

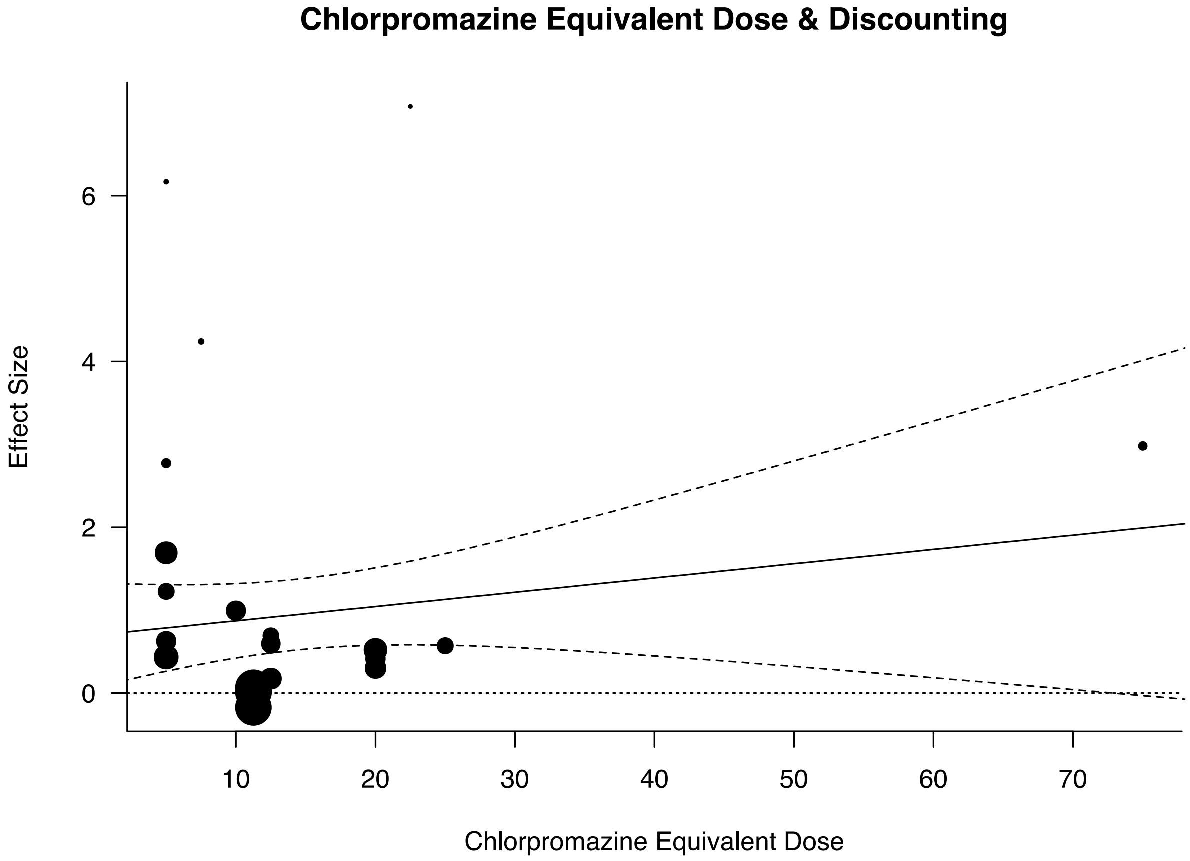

**Figure S11.** Meta-regression of placebo-controlled effect of D2:D3 receptor affinity ratio for D2-like agonists on reward discounting showing predicted regression slope and 95% confidence interval. Points represent individual effect sizes scaled by the ratio of the effect size to standard error. D2:D3 ratios smaller than 1 indicate greater affinity to D2 receptors and ratios larger than 1 indicate greater affinity to D3 receptors. Positive effect sizes indicate increased discounting on drug.

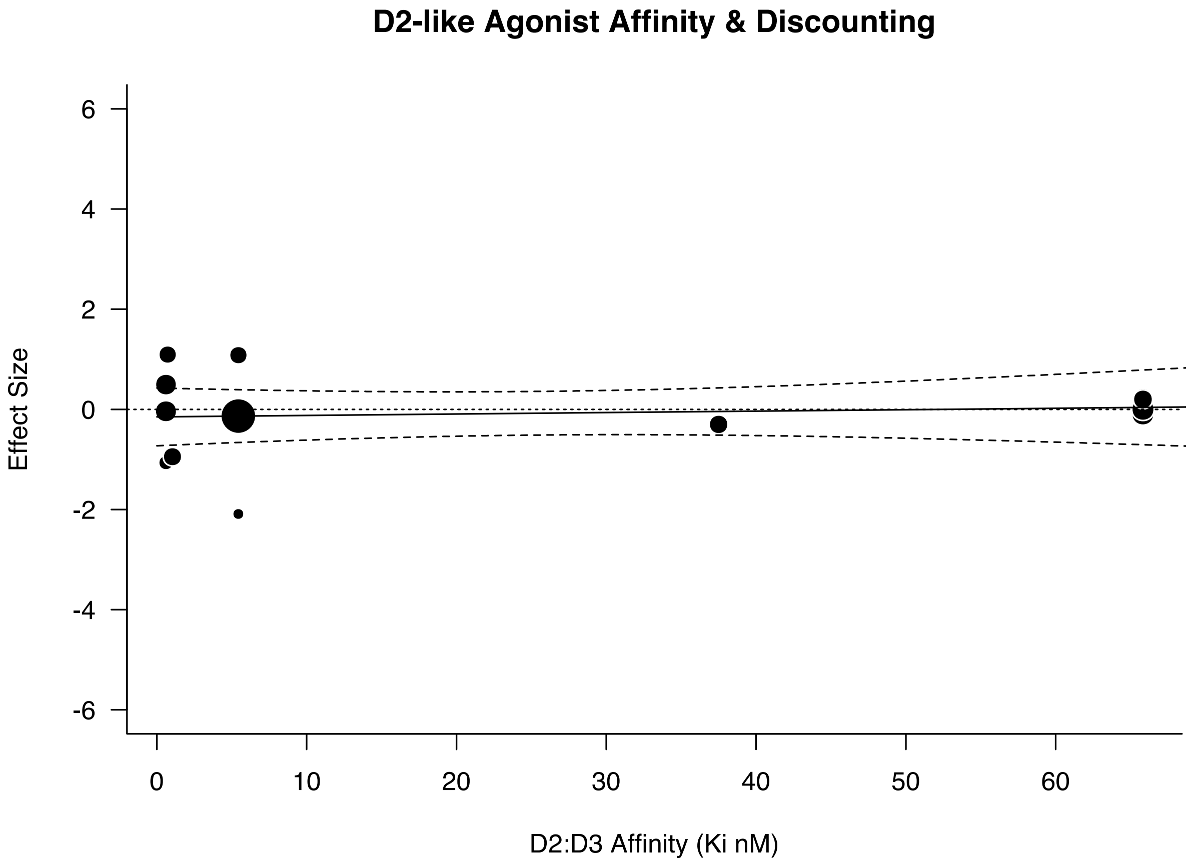

**Figure S12.** Meta-regression of placebo-controlled effect of D2:D3 receptor affinity ratio for D2-like antagonists on reward discounting showing predicted regression slope and 95% confidence interval. Points represent individual effect sizes scaled by the ratio of the effect size to standard error. D2:D3 ratios smaller than 1 indicate greater affinity to D2 receptors and ratios larger than 1 indicate greater affinity to D3 receptors. Positive effect sizes indicate increased discounting on drug.

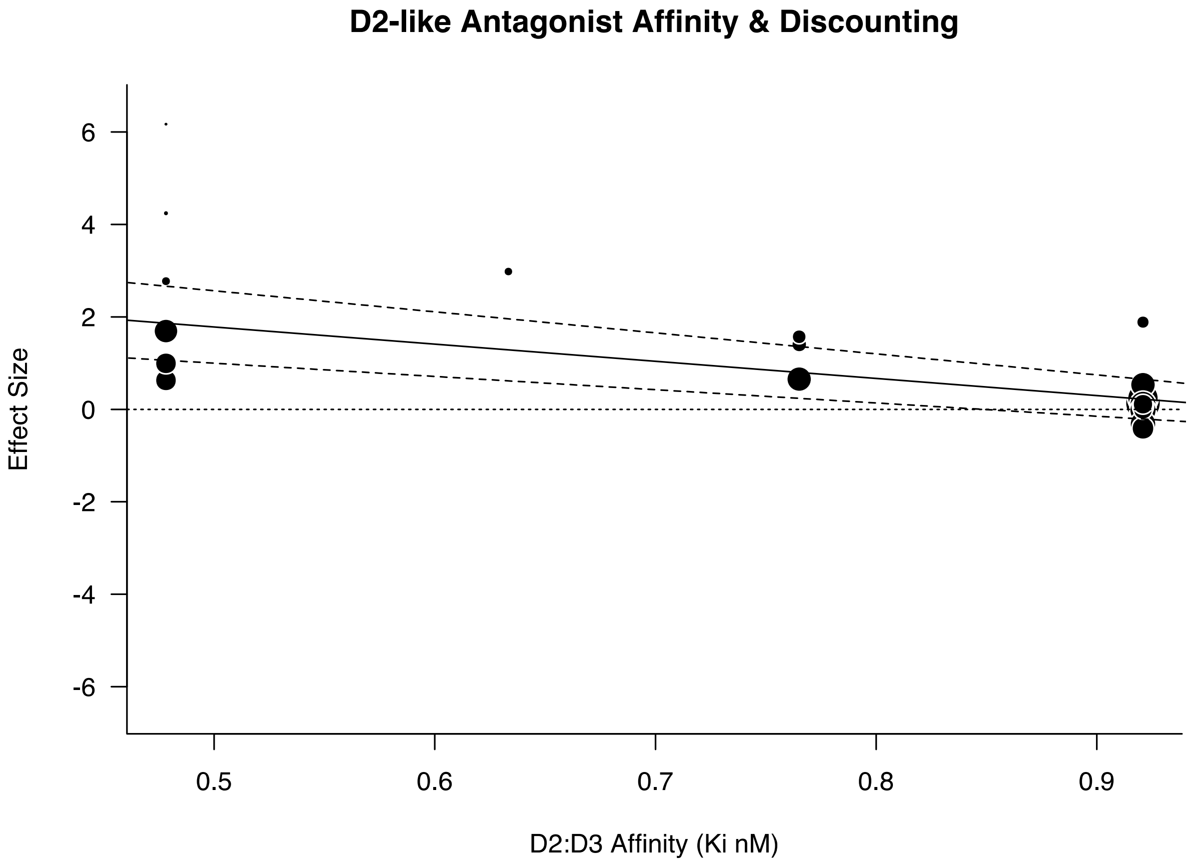

**Figure S13.** Histograms of D2-like agonist drugs sorted according to average identifiable D2:D3 affinity ratio (Ki nM). D2:D3 ratios smaller than 1 indicate greater affinity to D2 receptors and ratios larger than 1 indicate greater affinity to D3 receptors. Histograms include affinity ratio data prior to (left panel) and after (right panel) outlier removal.

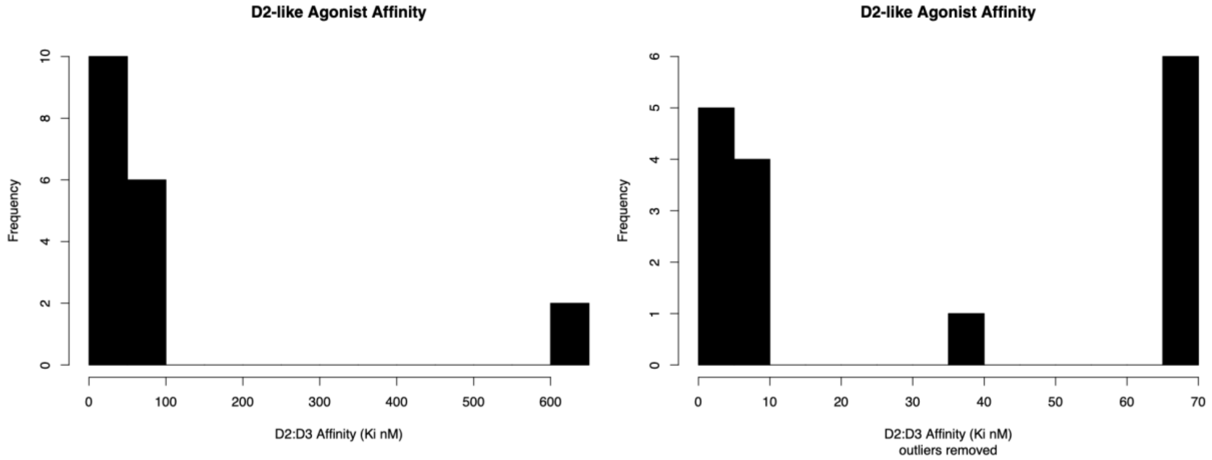


**Figure S14.** Histograms of D2-like agonist drugs sorted according to average identifiable D2:D3 affinity ratio (Ki nM). D2:D3 ratios smaller than 1 indicate greater affinity to D2 receptors and ratios larger than 1 indicate greater affinity to D3 receptors. Histograms include affinity ratio data prior to (left panel) and after (right panel) outlier removal.

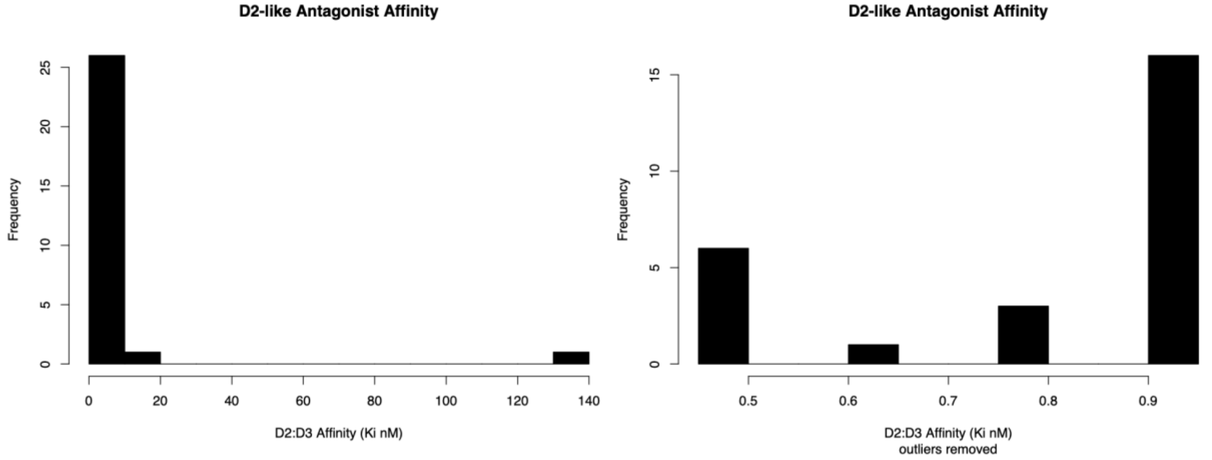

**Table S1.** Summary of extracted effect sizes for D1-like agonists. LE = Long Evans, LH = Lister Hooded, SD = Sprague-Dawley, W = Wistar. P = probability, T = time, propLarger = proportion of larger reward options chosen, MAD = mean adjusted delay.

| Author, Year | Rat Strain | Drug | Delivery | Dose | Cost Type | Discounting Measure | N | Effect Size | Standard Error |
|---|---|---|---|---|---|---|---|---|---|
| **St. Onge, 2009 [17]** | LE | SKF81297 | IP | 1 mg/kg | P | propLarger | 8 | -0.172 | 0.356 |
| **Koffarnus, 2011 [18]** | SD | SKF81297 | IP | 1 mg/kg | T | propLarger | 12 | 0.766 | 0.328 |
| **Yates, 2014 [5]** | SD | SKF81297 | mInf, mPFC | 0.4 μg | T | MAD | 6 | 0.719 | NA |
| | SD | SKF81297 | mInf, OFC | 0.4 μg | T | MAD | 7 | -0.181 | NA |
| **St. Onge, 2011 [6]** | LE | SKF81297 | mInf, mPFC | 0.4 μg | P | propLarger | 12 | -0.085 | 0.289 |
| **Larkin, 2016 [8]** | LE | SKF81297 | mInf, BLA | 1 μg | P | propLarger | 10 | 0.127 | 0.318 |
| | LE | SKF81297 | mInf, BLA | 1 μg | P | propLarger | 10 | 0.031 | 0.316 |
| **Stopper, 2013 [1]** | LE | SKF81297 | mInf, NAc | 2 μg/hemisphere | P | propLarger | 11 | 0.077 | 0.302 |
| **Simon, 2011 [19]** | LE | SKF81297 | IP | 1 mg/kg | P | propLarger | 12 | 0.224 | 0.292 |

**Table S2.** Summary of extracted effect sizes for D1-like antagonists. LE = Long Evans, LH = Lister Hooded, SD = Sprague-Dawley, W = Wistar. P = probability, T = time, E = effort, propLarger = proportion of larger reward options chosen, MAD = mean adjusted delay, IND PT = indifference point.

| Author, Year | Rat Strain | Drug | Delivery | Dose | Cost Type | Discounting Measure | N | Effect Size | Standard Error |
|---|---|---|---|---|---|---|---|---|---|

| | | | | | | | | | |
|---|---|---|---|---|---|---|---|---|---|
| **St. Onge, 2009 [17]** | LE | SCH23390 | IP | 0.01 mg/kg | P | propLarger | 8 | 0.39 | 0.367 |
| **Hosking, 2014 [20]** | LE | SCH23390 | IP | 0.01 mg/kg | E | propLarger | 28 | 0.033 | 0.189 |
| | LE | SCH23390 | IP | 0.01 mg/kg | E | propLarger | 22 | 0.22 | 0.216 |
| **Koffarnus, 2011 [18]** | SD | SCH23390 | IP | 0.032 mg/kg | T | propLarger | 12 | 0.572 | 0.311 |
| **Yates, 2014 [5]** | SD | SCH23390 | mInf, mPFC | 1 µg | T | MAD | 6 | 0.263 | NA |
| | SD | SCH23390 | mInf, OFC | 1 µg | T | MAD | 7 | -0.087 | NA |
| **Sink, 2007 [21]** | SD | SCH39166 | IP | 0.2 mg/kg | E | IND PT | 8 | 7.923 | 2.012 |
| **Cousins, 1994 [22]** | SD | SCH23390 | IP | 0.15 mg/kg | E | IND PT | 9 | 1.251 | 0.445 |
| **Li, 2015 [3]** | SD | SCH23390 | IP | 0.02 mg/kg | T | propLarger | 16 | 0.813 | 0.288 |
| **Wade, 2000 [23]** | SD | SCH23390 | IP | 20 µg/kg | T | IND PT | 17 | 0.035 | 0.243 |
| **St. Onge, 2011 [6]** | LE | SCH23390 | mInf, mPFC | 1 µg | P | propLarger | 12 | 0.247 | 0.293 |
| **Bardgett, 2009 [24]** | LE | SCH23390 | IP | 0.0125 mg/kg | E | IND PT | 9 | 2.482 | 0.673 |
| **Larkin, 2016 [8]** | LE | SCH23390 | mInf, BLA | 1 µg | P | propLarger | 8 | 0.311 | 0.362 |
| | LE | SCH23390 | mInf, BLA | 1 µg | P | propLarger | 8 | 0.316 | 0.362 |
| **Pattij, 2014 [9]** | W | SCH23390 | mInf, Insula | 1 µg/hemisphere | T | propLarger | 13 | 0.715 | 0.311 |
| **Stopper, 2013 [1]** | LE | SCH23390 | mInf, NAc | 1 µg/hemisphere | P | propLarger | 13 | 0.373 | 0.287 |
| **Pardey, 2013 [7]** | W | SCH23390 | mInf, mPFC | 3 µg/hemisphere | T | propLarger | 10 | 1.445 | 0.452 |

| | | | | | | | | | |
|---|---|---|---|---|---|---|---|---|---|
| | W | SCH23390 | mInf, OFC | 3 μg/hemisphere | T | propLarger | 10 | 1.098 | 0.4 |
| **Simon, 2011 [19]** | LE | SCH23390 | IP | 0.03 mg/kg | P | propLarger | 12 | 0.014 | 0.289 |

**Table S3.** Summary of extracted effect sizes for D2-like agonists. LE = Long Evans, LH = Lister Hooded, SD = Sprague-Dawley, W = Wistar. P = probability, T = time, E = effort, propLarger = proportion of larger reward options chosen, propSmaller = proportion of smaller reward options chosen, MAD = mean adjusted delay, IND PT = indifference point.

| Author, Year | Rat Strain | Drug | Delivery | Dose | Cost Type | Discounting Measure | N | Effect Size | Standard Error |
|---|---|---|---|---|---|---|---|---|---|
| **St. Onge, 2009 [17]** | LE | Bromocriptine | IP | 5 mg/kg | P | propLarger | 8 | -1.066 | 0.443 |
| | LE | PD 128,907 | IP | 0.5 mg/kg | P | propLarger | 8 | 0.805 | 0.407 |
| | LE | PD 168,077 | IP | 5 mg/kg | P | propLarger | 8 | 0.096 | 0.354 |
| **Madden, 2010 [25]** | W | Pramipexole | SC | 0.3 mg/kg | T | propSmaller | 9 | NA | NA |
| **Rokosik, 2012 [26]** | SD | Pramipexole | IP | 2 mg/kg | P | propLarger | 10 | -2.089 | 0.564 |
| **Koffarnus, 2011 [18]** | SD | Apomorphine | IP | 0.32 mg/kg | T | propLarger | 12 | 1.094 | 0.365 |
| | SD | Pramipexole | IP | 0.32 mg/kg | T | propLarger | 12 | 1.081 | 0.363 |
| | SD | Sumanirole | IP | 3.2 mg/kg | T | propLarger | 12 | 0.28 | 0.294 |
| | SD | ABT-724 | IP | 3.2 mg/kg | T | propLarger | 12 | 0.139 | 0.29 |
| **Yates, 2014 [5]** | SD | Quinpirole | mInf, mPFC | 5 μg | T | MAD | 6 | 0.374 | NA |
| | SD | Quinpirole | mInf, OFC | 5 μg | T | MAD | 7 | 0.369 | NA |
| **St. Onge, 2011 [6]** | LE | Quinpirole | mInf, mPFC | 10 μg | P | propLarger | 12 | -0.094 | 0.289 |

| | | | | | | | | | |
|---|---|---|---|---|---|---|---|---|---|
| **Bardgett, 2009 [24]** | LE | 7-OH-DPAT | IP | 0.3 mg/kg | E | IND PT | 9 | -0.301 | 0.341 |
| **Pes, 2017 [27]** | LE | Pramipexole | SC | 0.3 mg/kg | P | propLarger | 30 | -0.134 | 0.183 |
| **Tremblay, 2017 [28]** | LE | Ropinirole | SC | 5 mg/kg | P | propLarger | 12 | -0.946 | 0.347 |
| **Larkin, 2016 [8]** | LE | Quinpirole | mInf, BLA | 10 µg | P | propLarger | 9 | 0.199 | 0.337 |
| | LE | Quinpirole | mInf, BLA | 10 µg | P | propLarger | 9 | 0.228 | 0.338 |
| **Stopper, 2013 [1]** | LE | Quinpirole | mInf, NAc | 10 µg/hemisphere | P | propLarger | 12 | 0 | 0.289 |
| | LE | Bromocriptine | mInf, NAc | 10 µg/hemisphere | P | propLarger | 11 | -0.04 | 0.302 |
| | LE | PD 128,907 | mInf, NAc | 3 µg/hemisphere | P | propLarger | 11 | 0.315 | 0.309 |
| **Simon, 2011 [19]** | LE | Bromocriptine | IP | 5 mg/kg | P | propLarger | 12 | 0.494 | 0.306 |

**Table S4.** Summary of extracted effect sizes for D2-like antagonists. LE = Long Evans, LH = Lister Hooded, SD = Sprague-Dawley, W = Wistar. P = probability, T = time, E = effort, propLarger = proportion of larger reward options chosen, propSmaller = proportion of smaller reward options chosen, MAD = mean adjusted delay, IND PT = indifference point.

| Author, Year | Rat Strain | Drug | Delivery | Dose | Cost Type | Discounting Measure | N | Effect Size | Standard Error |
|---|---|---|---|---|---|---|---|---|---|
| **St. Onge, 2010 [29]** | LE | Flupenthixol | IP | 0.4 mg/kg | P | propLarger | 16 | 0.523 | 0.267 |
| | LE | Flupenthixol | IP | 0.4 mg/kg | P | propLarger | 11 | 0.414 | 0.314 |
| | LE | Flupenthixol | IP | 0.4 mg/kg | P | propLarger | 12 | 0.301 | 0.295 |
| | LE | Eticlopride | IP | 0.03 mg/kg | P | propLarger | 8 | 0.518 | 0.377 |

| | | | | | | | | | |
|---|---|---|---|---|---|---|---|---|---|
| **St. Onge, 2009 [17]** | LE | L-745,870 | IP | 5 mg/kg | P | propLarger | 8 | -0.124 | 0.355 |
| **van Gaalen, 2006 [30]** | W | Eticlopride | IP | 0.09 mg/kg | T | propLarger | 16 | 0.532 | 0.267 |
| **Hosking, 2014 [20]** | LE | Eticlopride | IP | 0.06 mg/kg | E | propLarger | 28 | 0.13 | 0.19 |
| | LE | Eticlopride | IP | 0.06 mg/kg | E | propLarger | 22 | 0.244 | 0.216 |
| **Koffarnus, 2011 [18]** | SD | Haloperidol | IP | 0.1 mg/kg | T | propLarger | 12 | 0.627 | 0.316 |
| | SD | L-741,626 | IP | 3.2 mg/kg | T | propLarger | 12 | 0.585 | 0.312 |
| | SD | PG01037 | IP | 56 mg/kg | T | propLarger | 12 | 0.251 | 0.293 |
| | SD | L-745,870 | IP | 3.2 mg/kg | T | propLarger | 12 | -0.136 | 0.29 |
| **Yates, 2014 [5]** | SD | Eticlopride | mInf, mPFC | 1 µg | T | MAD | 7 | 2.216 | NA |
| | SD | Eticlopride | mInf, OFC | 1 µg | T | MAD | 7 | 0.575 | NA |
| **Sink, 2007 [21]** | SD | Eticlopride | IP | 0.1 mg/kg | E | IND PT | 10 | 1.887 | 0.527 |
| **Cousins, 1994 [22]** | SD | Haloperidol | IP | 0.15 mg/kg | E | IND PT | 7 | 4.241 | 1.195 |
| | SD | Flupenthixol | IP | 0.45 mg/kg | E | IND PT | 7 | 7.077 | 1.929 |
| | SD | Sulpiride | IP | 150 mg/kg | E | IND PT | 10 | 2.981 | 0.738 |
| **Li, 2015 [3]** | SD | Eticlopride | IP | 0.09 mg/kg | T | propLarger | 16 | -0.316 | 0.256 |
| | SD | Eticlopride | mInf, NAc Core | 2 µg/hemisphere | T | propLarger | 16 | 0.108 | 0.251 |
| | SD | Eticlopride | mInf, BLA | 2 µg/hemisphere | T | propLarger | 16 | -0.024 | 0.25 |
| **Denk, 2005 [31]** | LH | Haloperidol | IP | 0.2 mg/kg | T | propLarger | 15 | 0.994 | 0.316 |
| **Wade, 2000 [23]** | SD | Flupenthixol | IP | 100 µg/kg | T | IND PT | 17 | 0.434 | 0.254 |
| | SD | Raclopride | IP | 120 µg/kg | T | IND PT | 17 | 0.653 | 0.267 |
| | LE | Flupenthixol | IP | 0.5 mg/kg | E | propLarger | 8 | 0.571 | 0.381 |

| | | | | | | | | | |
|---|---|---|---|---|---|---|---|---|---|
| **Floresco, 2008 [32]** | LE | Flupenthixol | IP | 0.25 mg/kg | E | propLarger | 8 | 0.693 | 0.394 |
| | LE | Flupenthixol | IP | 0.25 mg/kg | T | propLarger | 8 | 0.576 | 0.382 |
| **Mai, 2015 [33]** | LH | Flupenthixol | IP | 0.25 mg/kg | P | propLarger | 11 | 0.597 | 0.327 |
| **St. Onge, 2011 [6]** | LE | Eticlopride | mInf, mPFC | 1 µg | P | propLarger | 12 | -0.411 | 0.301 |
| **Bardgett, 2009 [24]** | LE | Haloperidol | IP | 0.1 mg/kg | E | IND PT | 9 | 6.168 | 1.492 |
| | LE | U-99194 | IP | 6.26 mg/kg | E | IND PT | 9 | 0.387 | 0.346 |
| **Yohn, 2017 [2]** | SD | Haloperidol | IP | 0.1 mg/kg | E | IND PT | 10 | 2.773 | 0.696 |
| **Larkin, 2016 [8]** | LE | Eticlopride | mInf, BLA | 1 µg | P | propLarger | 9 | 0.111 | 0.334 |
| | LE | Eticlopride | mInf, BLA | 1 µg | P | propLarger | 10 | 0.058 | 0.316 |
| **Pattij, 2014 [9]** | W | Eticlopride | mInf, Insula | 1 µg/hemisphere | T | propLarger | 13 | 0.161 | 0.279 |
| **Stopper, 2013 [1]** | LE | Eticlopride | mInf, NAc | 1 µg/hemisphere | P | propLarger | 8 | -0.012 | 0.354 |
| **Pardey, 2013 [7]** | W | Raclopride | mInf, mPFC | 6 µg/hemisphere | T | propLarger | 10 | 1.569 | 0.472 |
| | W | Raclopride | mInf, OFC | 6 µg/hemisphere | T | propLarger | 10 | 1.404 | 0.446 |
| **Ostlund, 2012 [34]** | SD | Flupenthixol | IP | 0.225 mg/kg | E | PropLarger | 36 | 0.012 | 0.167 |
| | SD | Flupenthixol | IP | 0.225 mg/kg | E | PropLarger | 36 | 0.064 | 0.167 |
| | SD | Flupenthixol | IP | 0.225 mg/kg | E | PropLarger | 36 | -0.173 | 0.168 |
| **Shafiei, 2012 [35]** | LE | Flupenthixol | IP | 0.25 mg/kg | E | PropLarger | 12 | 0.174 | 0.291 |
| **Randall, 2012 [36]** | SD | Haloperidol | IP | 0.1 mg/kg | E | IND PT | 32 | 1.693 | 0.276 |
| **Simon, 2011 [19]** | LE | Eticlopride | IP | 0.05 mg/kg | P | propLarger | 12 | -0.014 | 0.289 |

| | | | | | | | | | |
|---|---|---|---|---|---|---|---|---|---|
| **Olmstead, 2006 [37]** | LE | Flupenthixol | IP | 0.1 mg/kg | T | propSmaller | 12 | 1.226 | 0.382 |

**Table S5.** Summary of extracted effect sizes for DAT-modulating drugs. LE = Long Evans, LH = Lister Hooded, SD = Sprague-Dawley, W = Wistar. P = probability, T = time, E = effort, propLarger = proportion of larger reward options chosen, MAD = mean adjusted delay, IND PT = indifference point.

| Author, Year | Rat Strain | Drug | Delivery | Dose | Cost Type | Discounting Measure | N | Effect Size | Standard Error |
|---|---|---|---|---|---|---|---|---|---|
| **Orsini, 2017 [4]** | LE | d-Amphetamine | mInf, NAc | 20 μg/hemisphere | T | propLarger | 15 | -0.301 | 0.264 |
| | LE | d-Amphetamine | mInf, NAc | 20 μg/hemisphere | T | propLarger | 11 | 0.513 | 0.321 |
| **St. Onge, 2010 [29]** | LE | d-Amphetamine | IP | 0.5 mg/kg | P | propLarger | 16 | 0.371 | 0.258 |
| | LE | d-Amphetamine | IP | 0.5 mg/kg | P | propLarger | 15 | -0.291 | 0.264 |
| | LE | d-Amphetamine | IP | 0.5 mg/kg | P | propLarger | 12 | -0.143 | 0.29 |
| **St. Onge, 2009 [17]** | LE | d-Amphetamine | IP | 1 mg/kg | P | propLarger | 8 | -0.404 | 0.368 |
| **van Gaalen, 2006 [30]** | W | Methylphenidate | IP | 3 mg/kg | T | propLarger | 16 | -0.884 | 0.295 |
| **Baarendse, 2012 [38]** | LH | GBR 12909 | IP | 10 mg/kg | T | propLarger | 14 | -1.293 | 0.362 |
| | LH | d-Amphetamine | IP | 1 mg/kg | T | propLarger | 14 | -0.513 | 0.284 |
| **Koffarnus, 2011 [18]** | SD | d-Amphetamine | IP | 1 mg/kg | T | propLarger | 12 | 0.35 | 0.297 |
| | SD | GBR 12909 | IP | 10 mg/kg | T | propLarger | 12 | 0.137 | 0.29 |

| | | | | | | | | | |
|---|---|---|---|---|---|---|---|---|---|
| **Yates, 2014 [5]** | SD | Methylphenidate | mInf, mPFC | 100 µg | T | MAD | 6 | -0.748 | NA |
| | SD | Methylphenidate | mInf, OFC | 100 µg | T | MAD | 7 | -0.594 | NA |
| | SD | d-Amphetamine | mInf, mPFC | 4 µg | T | MAD | 6 | -0.326 | NA |
| | SD | d-Amphetamine | mInf, OFC | 4 µg | T | MAD | 6 | -0.292 | NA |
| | SD | Atomexetine | mInf, mPFC | 16 µg | T | MAD | 5 | -0.421 | NA |
| | SD | Atomexetine | mInf, OFC | 16 µg | T | MAD | 6 | -0.459 | NA |
| **Cousins, 1994 [22]** | SD | d-Amphetamine | IP | 3 mg/kg | E | IND PT | 7 | 2.014 | 0.658 |
| **Siemian, 2017 [39]** | SD | d-Methamphetamine | IP | 1 mg/kg | T | propLarger | 8 | 0.027 | 0.354 |
| **Li, 2015 [3]** | SD | Cocaine | IP | 20 mg/kg | T | propLarger | 16 | -1.503 | 0.365 |
| **Wade, 2000 [23]** | SD | d-Amphetamine | IP | 1 mg/kg | T | IND PT | 17 | -0.818 | 0.28 |
| **Floresco, 2008 [32]** | LE | d-Amphetamine | IP | 0.5 mg/kg | E | propLarger | 8 | 0.545 | 0.379 |
| | LE | d-Amphetamine | IP | 0.5 mg/kg | T | propLarger | 8 | -0.037 | 0.354 |
| **Mai, 2015 [33]** | LH | d-Amphetamine | IP | 0.5 mg/kg | P | propLarger | 11 | -1.144 | 0.388 |
| **Yohn, 2016 [2]** | SD | GBR12909 | mInf, NAc Core | 10 mg/kg | E | IND PT | 18 | -0.878 | 0.277 |
| | SD | Atomoxetine | IP | 1 mg/kg | E | IND PT | 8 | 2.31 | 0.677 |
| **Zeeb, 2016 [40]** | LE | d-Amphetamine | IP | 1.5 mg/kg | T | propLarger | 30 | 0.445 | 0.191 |
| **Mai, 2015 [41]** | LH | d-Amphetamine | IP | 2 mg/kg | P | propLarger | 10 | -1.119 | 0.403 |
| | LH | Cocaine | IP | 15 mg/kg | P | propLarger | 10 | -0.478 | 0.334 |

| | | | | | | | | | |
|---|---|---|---|---|---|---|---|---|---|
| **Randall, 2015 [42]** | SD | Bupropion | IP | 40 mg/kg | E | IND PT | 42 | -0.73 | 0.174 |
| **Hernandez, 2014 [43]** | LE | Cocaine | IP | 10 mg/kg | T | propLarger | 6 | 1.131 | 0.523 |
| **Wiskerke, 2011 [44]** | W | d-Amphetamine | IP | 0.5 mg/kg | T | propLarger | 14 | -1.165 | 0.346 |
| **Barbelivien, 2008 [45]** | LE | d-Amphetamine | IP | 1 mg/kg | T | propLarger | 31 | -0.1 | 0.18 |
| **Sommer, 2014 [46]** | SD | MRZ 9547 | IP | 100 mg/kg | E | IND PT | 10 | -4.234 | 0.867 |
| | SD | MRZ 9546 | IP | 100 mg/kg | E | IND PT | 10 | -1.132 | 0.489 |
| | SD | Modafinil | IP | 64 mg/kg | E | IND PT | 11 | 0.356 | 0.441 |
| | SD | Methylphenidate | IP | 10 mg/kg | E | IND PT | 11 | -1.472 | 0.504 |
| | SD | d-Amphetamine | IP | 1 mg/kg | E | IND PT | 10 | -0.93 | 0.476 |

### References


1. Stopper CM, Khayambashi S, Floresco SB. Receptor-Specific Modulation of Risk-Based Decision Making by Nucleus Accumbens Dopamine. Neuropsychopharmacology. 2013;38:715–728.
2. Yohn SE, Alberati D, Correa M, Salamone JD. Assessment of a glycine uptake inhibitor in animal models of effort-related choice behavior: implications for motivational dysfunctions. Psychopharmacology (Berl). 2017;234:1525–1534.
3. Li Y, Zuo Y, Yu P, Ping X, Cui C. Role of basolateral amygdala dopamine D2 receptors in impulsive choice in acute cocaine-treated rats. Behav Brain Res. 2015;287:187–195.
4. Orsini CA, Mitchell MR, Heshmati SC, Shimp KG, Spurrell MS, Bizon JL, et al. Effects of nucleus accumbens amphetamine administration on performance in a delay discounting task. Behav Brain Res. 2017;321:130–136.
5. Yates JR, Perry JL, Meyer AC, Gipson CD, Charnigo R, Bardo MT. Role of medial prefrontal and orbitofrontal monoamine transporters and receptors in performance in an adjusting delay discounting procedure. Brain Res. 2014;1574:26–36.
6. St. Onge JR, Abhari H, Floresco SB. Dissociable Contributions by Prefrontal D1 and D2 Receptors to Risk-Based Decision Making. J Neurosci. 2011;31:8625–8633.
7. Pardey MC, Kumar NN, Goodchild AK, Cornish JL. Catecholamine receptors differentially mediate impulsive choice in the medial prefrontal and orbitofrontal cortex. J Psychopharmacol (Oxf). 2013;27:203–212.
8. Larkin JD, Jenni NL, Floresco SB. Modulation of risk/reward decision making by dopaminergic transmission within the basolateral amygdala. Psychopharmacology (Berl). 2016;233:121–136.
9. Pattij T, Schetters D, Schoffelmeer ANM. Dopaminergic modulation of impulsive decision making in the rat insular cortex. Behav Brain Res. 2014;270:118–124.
10. Cervantes-Arriaga A, Rodríguez-Violante M, Villar-Velarde A, Corona T. Cálculo de unidades de equivalencia de levodopa en enfermedad de Parkinson. Arch Neurocienc. 2009;14:116–119.
11. Gardner DM, Murphy AL, O’Donnell H, Centorrino F, Baldessarini RJ. International Consensus Study of Antipsychotic Dosing. Am J Psychiatry. 2010;167:686–693.
12. Leucht S, Samara M, Heres S, Patel MX, Woods SW, Davis JM. Dose Equivalents for Second-Generation Antipsychotics: The Minimum Effective Dose Method. Schizophr Bull. 2014;40:314–326.
13. Smith C. Levodopa dose equivalency: A systematic review2010.
14. Woods SW. Chlorpromazine equivalent doses for the newer atypical antipsychotics. J Clin Psychiatry. 2003;64:663–667.
15. Landwehrmeyer B, Mengod G, Palacios JM. Differential Visualization of Dopamine D2 and D3 Receptor Sites in Rat Brain. A Comparative Study Using *In Situ* Hybridization Histochemistry and Ligand Binding Autoradiography. Eur J Neurosci. 1993;5:145–153.
16. Soto PL, Hiranita T, Xu M, Hursh SR, Grandy DK, Katz JL. Dopamine D2-Like Receptors and Behavioral Economics of Food Reinforcement. Neuropsychopharmacology. 2016;41:971–978.
17. St Onge JR, Floresco SB. Dopaminergic Modulation of Risk-Based Decision Making. Neuropsychopharmacology. 2009;34:681–697.

18. Koffarnus MN, Newman AH, Grundt P, Rice KC, Woods JH. Effects of selective dopaminergic compounds on a delay-discounting task: Behav Pharmacol. 2011;22:300–311.
19. Simon NW, Montgomery KS, Beas BS, Mitchell MR, LaSarge CL, Mendez IA, et al. Dopaminergic Modulation of Risky Decision-Making. J Neurosci. 2011;31:17460–17470.
20. Hosking JG, Floresco SB, Winstanley CA. Dopamine Antagonism Decreases Willingness to Expend Physical, But Not Cognitive, Effort: A Comparison of Two Rodent Cost/Benefit Decision-Making Tasks. Neuropsychopharmacology. 2015;40:1005–1015.
21. Sink KS, Vemuri VK, Olszewska T, Makriyannis A, Salamone JD. Cannabinoid CB1 antagonists and dopamine antagonists produce different effects on a task involving response allocation and effort-related choice in food-seeking behavior. Psychopharmacology (Berl). 2008;196:565–574.
22. Cousins MS, Wei W, Salamone JD. Pharmacological characterization of performance on a concurrent lever pressing/feeding choice procedure: effects of dopamine antagonist, cholinomimetic, sedative and stimulant drugs. Psychopharmacology (Berl). 1994;116:529–537.
23. Wade TR, de Wit H, Richards JB. Effects of dopaminergic drugs on delayed reward as a measure of impulsive behavior in rats. Psychopharmacology (Berl). 2000;150:90–101.
24. Bardgett ME, Depenbrock M, Downs N, Points M, Green L. Dopamine modulates effort-based decision making in rats. Behav Neurosci. 2009;123:242–251.
25. Madden GJ, Johnson PS, Brewer AT, Pinkston JW, Fowler SC. Effects of pramipexole on impulsive choice in male wistar rats. Exp Clin Psychopharmacol. 2010;18:267–276.
26. Rokosik SL, Napier TC. Pramipexole-Induced Increased Probabilistic Discounting: Comparison Between a Rodent Model of Parkinson's Disease and Controls. Neuropsychopharmacology. 2012;37:1397–1408.
27. Pes R, Godar SC, Fox AT, Burgeno LM, Strathman HJ, Jarmolowicz DP, et al. Pramipexole enhances disadvantageous decision-making: Lack of relation to changes in phasic dopamine release. Neuropharmacology. 2017;114:77–87.
28. Tremblay M, Silveira MM, Kaur S, Hosking JG, Adams WK, Baunez C, et al. Chronic D2/3 agonist ropinirole treatment increases preference for uncertainty in rats regardless of baseline choice patterns. Eur J Neurosci. 2017;45:159–166.
29. St. Onge JR, Chiu YC, Floresco SB. Differential effects of dopaminergic manipulations on risky choice. Psychopharmacology (Berl). 2010;211:209–221.
30. van Gaalen MM, van Koten R, Schoffelmeer ANM, Vanderschuren LJMJ. Critical Involvement of Dopaminergic Neurotransmission in Impulsive Decision Making. Biol Psychiatry. 2006;60:66–73.
31. Denk F, Walton ME, Jennings KA, Sharp T, Rushworth MFS, Bannerman DM. Differential involvement of serotonin and dopamine systems in cost-benefit decisions about delay or effort. Psychopharmacology (Berl). 2005;179:587–596.
32. Floresco SB, Tse MTL, Ghods-Sharifi S. Dopaminergic and Glutamatergic Regulation of Effort- and Delay-Based Decision Making. Neuropsychopharmacology. 2008;33:1966–1979.
33. Mai B, Sommer S, Hauber W. Dopamine D1/D2 Receptor Activity in the Nucleus Accumbens Core But Not in the Nucleus Accumbens Shell and Orbitofrontal Cortex Modulates Risk-Based Decision Making. Int J Neuropsychopharmacol. 2015;18:pyv043.
34. Ostlund SB, Kosheleff AR, Maidment NT. Relative Response Cost Determines the Sensitivity of Instrumental Reward Seeking to Dopamine Receptor Blockade. Neuropsychopharmacology. 2012;37:2653–2660.

35. Shafiei N, Gray M, Viau V, Floresco SB. Acute Stress Induces Selective Alterations in Cost/Benefit Decision-Making. Neuropsychopharmacology. 2012;37:2194–2209.
36. Randall PA, Pardo M, Nunes EJ, López Cruz L, Vemuri VK, Makriyannis A, et al. Dopaminergic Modulation of Effort-Related Choice Behavior as Assessed by a Progressive Ratio Chow Feeding Choice Task: Pharmacological Studies and the Role of Individual Differences. PLoS ONE. 2012;7:e47934.
37. Olmstead MC, Hellemans KGC, Paine TA. Alcohol-induced impulsivity in rats: an effect of cue salience? Psychopharmacology (Berl). 2006;184:221–228.
38. Baarendse PJJ, Vanderschuren LJMJ. Dissociable effects of monoamine reuptake inhibitors on distinct forms of impulsive behavior in rats. Psychopharmacology (Berl). 2012;219:313–326.
39. Siemian JN, Xue Z, Blough BE, Li J-X. Comparison of some behavioral effects of d- and l-methamphetamine in adult male rats. Psychopharmacology (Berl). 2017;234:2167–2176.
40. Zeeb FD, Soko AD, Ji X, Fletcher PJ. Low Impulsive Action, but not Impulsive Choice, Predicts Greater Conditioned Reinforcer Salience and Augmented Nucleus Accumbens Dopamine Release. Neuropsychopharmacology. 2016;41:2091–2100.
41. Mai B, Hauber W. Orbitofrontal or accumbens dopamine depletion does not affect risk-based decision making in rats. Cogn Affect Behav Neurosci. 2015;15:507–522.
42. Randall PA, Lee CA, Podurgiel SJ, Hart E, Yohn SE, Jones M, et al. Bupropion Increases Selection of High Effort Activity in Rats Tested on a Progressive Ratio/Chow Feeding Choice Procedure: Implications for Treatment of Effort-Related Motivational Symptoms. Int J Neuropsychopharmacol. 2015;18:pyu017–pyu017.
43. Hernandez G, Oleson EB, Gentry RN, Abbas Z, Bernstein DL, Arvanitogiannis A, et al. Endocannabinoids Promote Cocaine-Induced Impulsivity and Its Rapid Dopaminergic Correlates. Biol Psychiatry. 2014;75:487–498.
44. Wiskerke J, Schetters D, van Es IE, van Mourik Y, den Hollander BRO, Schoffelmeer ANM, et al. -Opioid Receptors in the Nucleus Accumbens Shell Region Mediate the Effects of Amphetamine on Inhibitory Control But Not Impulsive Choice. J Neurosci. 2011;31:262–272.
45. Barbelivien A, Billy E, Lazarus C, Kelche C, Majchrzak M. Rats with different profiles of impulsive choice behavior exhibit differences in responses to caffeine and d-amphetamine and in medial prefrontal cortex 5-HT utilization. Behav Brain Res. 2008;187:273–283.
46. Sommer S, Danysz W, Russ H, Valastro B, Flik G, Hauber W. The dopamine reuptake inhibitor MRZ-9547 increases progressive ratio responding in rats. Int J Neuropsychopharmacol. 2014;17:2045–2056.
